# Unclusterable, underdispersed arrangement of insect-pollinated plants in pollinator niche space

**DOI:** 10.1101/2020.03.19.999169

**Authors:** Carlos M. Herrera

## Abstract

Pollinators can mediate facilitative or competitive relationships between plant species, but the comparative importance of these two conflicting phenomena in shaping community-wide pollinator resource use remains unexplored. This paper examines the idea that the arrangement in pollinator niche space of plant species samples comprising complete or nearly complete regional or local plant communities can help to evaluate the relative importance of facilitation and competition as drivers of community-wide pollinator resource use. Pollinator composition data for insect-pollinated plants from the Sierra de Cazorla mountains (southeastern Spain), comprising 85% of families and ~95% of widely distributed insect-pollinated species, were used to address the following questions at regional (45 sites, 221 plant species) and local (one site, 73 plant species) spatial scales: (1) Do objectively identifiable plant species clusters occur in pollinator niche space ? Four different pollinator niche spaces were considered whose axes were defined by insect orders, families, genera and species; and (2) If all plant species form a single, indivisible cluster in pollinator niche space, Are they overdispersed or underdispersed relative to a random arrangement ? “Clusterability” tests failed to reject the null hypothesis that there was only one pollinator-defined plant species cluster in pollinator niche space, irrespective of spatial scale, pollinator niche space or pollinator importance measurement (proportions of pollinator individuals or flowers visited by each pollinator type). Observed means of pairwise interspecific dissimilarity in pollinator composition were smaller than randomly simulated values in the order-, family- and genus-defined pollinator niche spaces at both spatial scales, thus revealing significantly non-random, underdispersed arrangement of plant species within the single cluster existing in each of these pollinator niche spaces. In the undisturbed montane habitats studied, arrangement of insect-pollinated plant species in pollinator niche space did not support a major role for interspecific competition as a force shaping community-wide pollinator resource use by plants, but rather suggested a situation closer to the facilitation-dominated extreme in a hypothetical competition-facilitation gradient. Results also highlight the importance of investigations on complete or nearly complete insect-pollinated plant communities for addressing novel hypotheses on the ecology and evolution of plant-pollinator systems.

## Introduction

The maturation of a full-fledged niche theory was a significant landmark in the trajectory of twentieth-century ecology (Vandermeer 1972, Chase and Leibold 2003, Ricklefs 2012). originally aimed at investigating how many and how similar coexisting species could be in a community, niche theory was predominantly concerned with animal communities and rested on the overarching assumption that interspecific competition for limiting resources is a major driving force in shaping community composition and resource use by individual species (Chase and Leibold 2003). Notions derived from niche theory sometimes also made their way into plant community studies. In particular, the niche theory framework is behind investigations of plant-pollinator interactions that have envisaged animal pollinators as “resources” exploited by insect-pollinated plants to ensure sexual reproduction, since competition for pollinators may result in loss of pollinator visits (exploitation) and/or disruption of conspecific pollen flow (interference) (Waser 1978, Pleasants 1980). In this context, pollinator partitioning over daily or seasonal gradients (Herrera 1990, Stone et al. 1998, Aizen and Vázquez 2006, de Souza et al. 2017), generalization *vs*. specialization on pollinators as alternative strategies of resource use (Herrera 1996, 2005, Waser et al. 1996, Armbruster 2017), divergence in pollinator composition as a mechanism alleviating interspecific competition (*Moldenke* 1975, Kephart 1983, Johnson 2010, Bouman et al. 2016, Johnson and Bronstein 2019), or pollinator-mediated plant species coexistence (Paw 2013, Benadi 2015, Benadi and Paw 2018), are all notions rooted in niche theory that to a greater or lesser extent share the canonical assumption that competition is an important driving force.

One insufficiently acknowledged limitation of the application of competition-laden niche theory to plant-pollinator systems is that, under certain circumstances, pollinator sharing by different plant species can result in facilitation rather than competition (Thomson 1978, Rathcke 1983, Callaway 1995). This possibility has been verified experimentally many times (see Braun and Lortie 2019 for review). Proximate mechanisms leading to pollinator-mediated interspecific facilitation include the “magnet species effect”, whereby plant species highly attractive to pollinators can benefit co-flowering ones in the neighborhood that are visited less often (Laverty 1992, Johnson et al. 2003, Hansen et al. 2007, Molina-Montenegro et al. 2008, Pellegrino et al. 2016); sequential facilitation, whereby earlier-flowering plant species benefit later-flowering ones at the same location by sustaining populations of site-faithful, shared pollinators (Waser and Real 1979, Ogilvie and Thomson 2016); floral mimicry, in which plant species can derive reproductive success when the colour, shape or scent of their flowers resemble those of a co-blooming plant (Gumbert and Kunze 2001, Johnson et al. 2003, Pellegrino et al. 2008); and diffuse beneficial effects of floral diversity at the plant community level on the pollination success of individual species (Ghazoul 2006, Seifan et al. 2014, Norfolk et al. 2016).

While the reality of both facilitative and competitive phenomena in plant-pollinator systems is well established, their comparative importance as drivers of pollinator resource use in natural communities remain essentially unexplored to date (but see, e.g., Moeller 2004, Tur et al. 2016, Eisen et al. 2019). The deleterious effect of interspecific pollen transfer due to pollinator sharing has been traditionally interpreted as supporting a role of competition in organizing plant-pollinator interactions by favoring pollinator partitioning (Stebbins 1970, Waser 1978, Wilson and Thomson 1996, Morales and Traveset 2008, Mitchell et al. 2009), yet recent studies suggest that some plant species can tolerate high levels of heterospecific pollen transfer without major detrimental effects on reproduction (Arceo-Gómez et al. 2018, Fang et al. 2019, Ashman et al. 2020). The hitherto little-explored possibility thus remains that facilitative interactions among animal-pollinated plants could also be significant drivers of plant-pollinator interactions at the community level, since the net outcome of opposing facilitative and competitive forces will depend on environmental context (Moeller 2004, Ghazoul 2006, Ye et al. 2014, Fitch 2017, Eisen et al. 2019) and the benefits of sharing pollinators can sometimes outweigh its costs (Tur et al. 2016). I propose here that the comparative importance of competition *vs*. facilitation in shaping community-wide plant-pollinator systems could be evaluated by examining the arrangement of plant species in the pollinator niche space, as outlined in the next section.

### A testable hypothesis on species arrangement in pollinator niche space

Pollinator-mediated interspecific facilitation and competition can be envisaged as exerting opposite forces on pollinator resource use, tending to favor pollinator sharing and divergence, respectively. Insights into the relative importance of these two conflicting forces could thus be gained by examining how *the whole set* of animal-pollinated species in a plant community are arranged in the multidimensional niche space whose axes are biologically distinct pollinator “types”. Such types should include all objectively recognizable pollinator classes that are sufficiently different in morphology, behavior and, in general, relevant pollinating features as to represent distinct resources (i.e., niche axes) from the plants’ perspective. A simple sketch of this idea referred to just two pollinator niche axes is shown in Fig. 1. Divergence in pollinator resource use, denoted by two or more plant species clusters in pollinator space (Fig. 1A), would be consistent with competition playing a major role in the community-level organization of plant-pollinator interactions. This assumes that divergence would be favored whenever there are some deleterious effects derived from sharing the same pollinators, irrespective of whether competition is by exploitation (reduction in pollinator visitation) or interference (disruption of pollen transfer). A single cluster in the pollinator niche space (Figs. 1B and 1C) would be suggestive of predominant facilitation having promoted convergence in pollinator resource use. The arrangement of plant species within individual clusters would also be informative. Densely packed species (underdispersed relative to a random distribution, Fig 1C) would support the interpretation that facilitation has played a major role, while loosely packed species (overdispersed relative to a random distribution, Fig. 1B) would instead be suggestive of competition having favored repulsion in niche space via limiting similarity effects. It is thus proposed that the importance of interspecific competition for pollinators as a major driving force shaping the arrangement of plants in the pollinator niche space should decline from Fig. 1A to 1C. Since variation in important pollinating features (e.g., frequency of pollen transfer, size and quality of pollen loads delivered, location of pollen grains on body surfaces) is generally related to the affiliation at various levels of the taxonomic hierarchy, taxonomic categories provide straightforward, objectively-defined proxies for relevant resource axes in pollinator niche space. This admittedly simplified approach will be adopted in this paper, although it must be acknowledged that pollinators in different taxonomic groups can be similar in pollinating features (Herrera 1987), the same taxonomic entity can include heterogeneous pollinators (e.g., sexes of the same species; Tang et al. 2019), and plant species could also partition pollinators by placing pollen on different parts of their bodies (Muchhala and Thomson 2012).

**Fig. 1.**
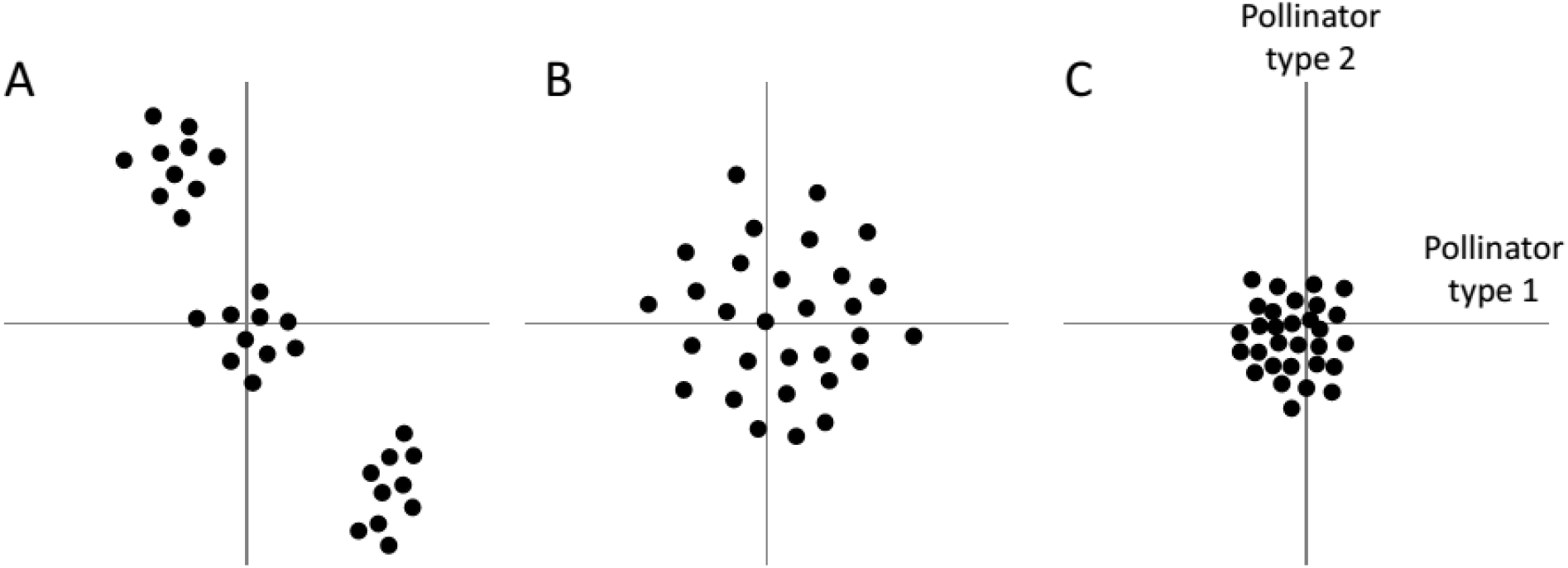
Schematic depiction of three possible arrangements of 30 species of animal-pollinated plants (black dots) in an imaginary pollinator niche space defined by two pollinator types (horizontal and vertical axes). The coordinates of each plant species on the axes are the relative contributions of pollinator types to its pollinator assemblage, scaled and centered. It is proposed that the importance of interspecific competition for pollinators as a major driving force shaping the arrangement of plants in the pollinator niche space should decline from A to C (see text).

The main objective of this paper is to provide a case example for the preceding framework by examining the arrangement in the pollinator niche space of a very large sample of insect-pollinated species from a well-preseved montane plant community. A secondary objective is to highlight the importance of adopting “zooming out” approaches in the study of the ecology of plant-pollinator interactions (Herrera 2020). Current knowledge of pollination system niches has been mostly obtained from studies dealing with small species samples (usually < 20 species), or focusing on specific pollination guilds of closely related plants associated with particular pollinators. While irreplaceable for other purposes, these investigations lack the biological amplitude needed to address broad-scale questions and warrant generalizations on the overall organization of plant-pollinator systems (Johnson 2010, Herrera 2019, 2020). Only a comprehensive coverage of the ecological and phylogenetic range of the insect-pollinated plants in a region and their whole set of pollinators could help to discriminate among the scenarios hypothesized in Fig. 1. The species sample examined here virtually encompasses the whole community of insect-pollinated plants in a southeastern Spanish biodiversity hotspot (Sierra de Cazorla mountains; see Molina-Venegas et al. 2015, and *Study area and plant species sample* below) and the whole spectrum of their insect pollinators.

The following two specific questions will be addressed: (1) Are there distinct, objectively identifiable clusters of plant species in the pollinator niche space ? Four different pollinator niche spaces with broadly different dimensionalities will be considered, with axes defined by insect orders, families, genera and species; and (2) If plant species form a single indivisible cluster in a given pollinator niche space, Is there evidence for either overdispersed or underdispersed arrangement within the cluster ? Two simplifications in the scheme of Fig. 1 must be acknowledged. First, the sign and/or magnitude of the hypothesized competition-facilitation balance mediated by pollinators may depend on the spatial scale considered (Tur et al. 2016, Braun and Lortie 2019, Eisen et al. 2019). And second, the scheme does not incorporate the possibility that even if divergence in pollinator composition could not be detected, pollinator partitioning associated with differential blooming time or habitat segregation could still reflect the effect of interspecific competition. To account for these potentially confounding factors, questions (1) and (2) will be addressed at two different spatial scales by conducting separate analyses for the whole set of plant species sampled in the region (“regional scale” hereafter) and for a large subset of them that coexist locally in an intensively studied plant community (“local scale”). In addition, the possible relationships between interspecific differences in habitat type and divergence in flowering time, and interspecific divergence in pollinator composition, will also be addressed.

## Materials and Methods

### Study area and plant species sample

Pollinator composition data considered in this paper were collected during February-December 1997-2019 (with a gap in 2000-2002) at 45 sites in the Sierras de Cazorla-Segura-Las Villas Natural Park (Jaén Province, southeastern Spain), a region characterized by outstanding biological diversity and well-preserved mountain habitats. The geographical distribution of sampling sites is shown in Fig. 2. Quantitative data on pollinator composition were obtained for 221 plant species in 48 families (Appendix S1: Table S1). This sample comprises 85% of families and ~95% of widely distributed species of insect-pollinated plants in the study area (Gómez Mercado 2011, C. M. Herrera, *unpublished data*), and is a superset of the 191 species in 43 families studied by Herrera (2020: Table 1). Asteraceae (41 species), Lamiaceae (24), Fabaceae (14), Brassicaceae (12), Rosaceae (11) and Ranunculaceae (9) contributed most species to the sample. About two thirds of species (*N* = 156) were sampled for pollinators only one year, and the rest were sampled ≥ 2 years. In the vast majority of species (*N* = 211) pollinator sampling was conducted on a single site. Pollinators of some species considered here vary among sites or years, but such intraspecific variation is quantitatively minor in comparison to the broad interspecific range occurring in the large species sample considered here, as previously shown by Herrera (2020: Table 2, Fig. 2) by means of formal variance partitions. Data for the same species obtained in more than one site or year were thus combined into a single sample for the analyses. For one third of species (*N* = 73; Appendix S1: Table S1) pollinator data were obtained in an area of ~10 ha at the intensively studied Nava de las Correhuelas site (Fig. 2). These species represent ~90% of all insect-pollinated species known to coexist locally at the site, and will form the basis for analyses at the local scale.

**Fig. 2.**
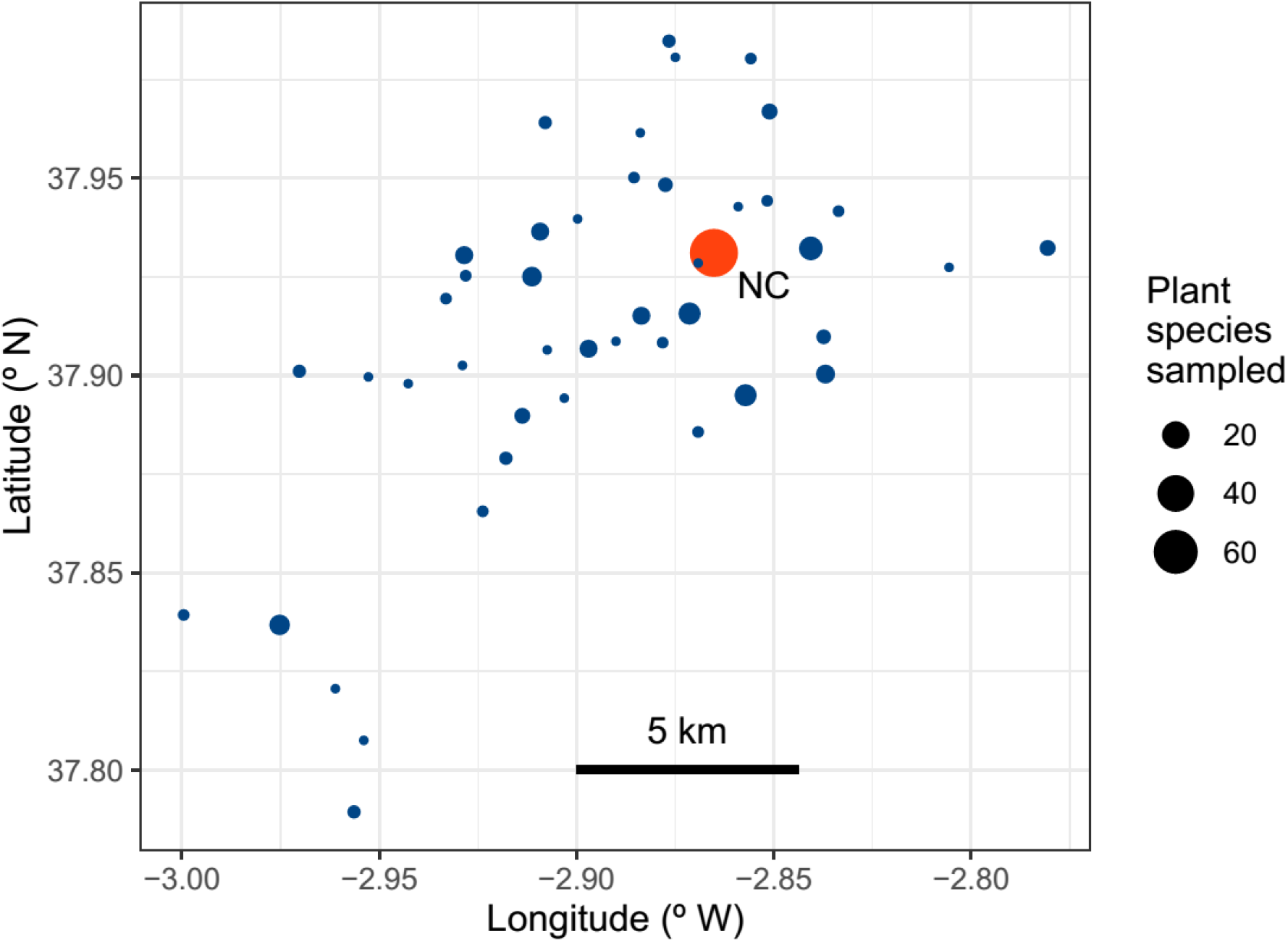
Map showing the location in the Sierra de Cazorla region, southeastern Spain, of the *N* = 45 sites where pollinator censuses were carried out. The area of each dot is proportional to the number of plant species sampled at the site. The large red dot is the intensively studied Nava de las Correhuelas site (NC, *N* = 73 plant species sampled), which provided the data used in the analyses at the local scale.

**Table 1.** Pollinator niche space dimensionality and sample sizes for the different combinations of spatial scale and niche space considered in this study, all plant species combined.

| Spatial scale | Pollinator niche axes <sup>1</sup> | Pollinator individuals | Flower visits |
| --- | --- | --- | --- |
| Regional (45 sites, 221 plant species) |  |  |  |
| | Orders ($n = 7$) | 33,835 | 107,626 |
| | Families ($n = 94$) | 33,835 | 107,626 |
| | Genera ($n = 346$) | 33,245 | 106,630 |
| | Species ($n = 542$) | 27,201 | 93,401 |
| Local (1 site, 73 plant species) |  |  |  |
| | Orders ($n = 7$) | 13,014 | 36,492 |
| | Families ($n = 78$) | 13,014 | 36,492 |
| | Genera ($n = 266$) | 12,755 | 36,059 |
| | Species ($n = 386$) | 10,237 | 30,838 |

*Notes:* For a given pollinator niche space, only pollinator visitation records where the pollinator was identified at the corresponding taxonomic resolution were considered in the analyses.
<sup>1</sup> $n$ = number of dimensions of the corresponding niche space.

Pollinator sampling dates for each species roughly matched its peak flowering date. The mean date of pollinator censuses for each species, expressed as days from 1 January, was used in analyses relating differences in pollinator composition to variation in flowering phenology. Mean census dates for the species studied ranged between 39–364 days from 1 January, and most species flowered during May-July (interquartile range = 145-190 days from 1 January) (Appendix S1: Table S1). For analyses relating differences in pollinator composition to dissimilarity in habitat type, each plant species was classed into one of the following nine major habitat categories (total species per habitat in parentheses; see also Herrera 2020): banks of permanent streams or flooded/damp areas around springs (18); dwarf mountain scrub dominated by cushion plants (27); forest edges and large clearings (34); forest interior (20); patches of grasslands and meadows on deep soils in relatively flat terrain (40); local disturbances caused by humans, large mammals or natural abiotic processes (24); sandy or rocky dolomitic outcrops (29); tall, dense Mediterranean sclerophyllous forest and scrub (17); vertical rock cliffs (12 species). Individual species’ assignments to habitat types are shown in Appendix S1: Table S1.

### Pollinator composition

Data on pollinator composition were obtained for every species as described in Herrera (2019, 2020). The elemental sampling unit was the “pollinator census”, consisting of a 3-min watch of a flowering patch whose total number of open flowers was also counted. All pollinators visiting some flower in the focal patch during the 3-min period were identified, and total number of flowers probed by each individual was recorded. Detailed information on sampling effort for individual plant species is shown in Appendix S1: Table S1. For all plant species combined, a total of 28,128 censuses were conducted on 575 different dates. Assessing the number of flowers visited by pollinators was impractical in the case of species with tiny flowers densely packed into compact inflorescences (e.g., Apiaceae, Asteraceae). In these species the number of inflorescences available per patch and visited per census were counted rather than individual flowers, although for simplicity I will always refer to “flower visitation”. A total of approximately 135,000 close-up photographs of insects visiting flowers were routinely taken during censuses using a DSLR digital camera and 105 mm macro lens, which were used for insect identification, keeping photographic vouchers, and ascertaining the pollinating status of different insect taxa. Based on these photographs, only taxa whose individuals contacted anthers or stigmas, or with discernible pollen grains on their bodies, were considered as pollinators and included in this study. This approach is admittedly crude, as it ignores possible differences among pollinators in pollinating effectiveness, but it is the only feasible one when dealing with many species of plants and pollinators. Despite its limitations, however, my photographically-informed method is superior and more biologically realistic than the assignment of any floral visitor to the “pollinator” category often used in recent plant-pollinator community studies.

### Data analysis

Four different pollinator niche spaces and two measurements of pollinator importance were considered throughout the analyses, each of which was carried out separately at the regional and local scales. Sample characteristics for every analysis are summarized in Table 1. Pollinator composition for each plant species was assessed using the proportion of pollinator individuals recorded and the proportion of flowers visited for each pollinator type, which are termed “proportion of visitors” and “proportion of visits”, respectively. Four distinct, taxonomically-defined classifications of pollinators were considered, based respectively on their affiliation to insect orders, families, genera and species. These classifications defined four pollinator niche spaces that differed in dimensionality and the biological meaning of their axes (Table 1). For a given pollinator niche space, only pollinator visitation records where the pollinator had been identified at the corresponding taxonomic resolution were included in the analyses. Out of a total of 33,835 individual pollinators recorded in censuses, 100% were identified to order and family, 98.3% to genus, and 80.3% to species (i.e., a full Latin binomial), which explains variation among niche spaces in sample characteristics (Table 1).

The question of whether plant species clusters could be recognized in each of the four pollinator niche spaces considered was first addressed informally by comparing the distribution of plant species in a reduced bidimensional space with the various scenarios envisaged in Fig. 1. For each combination of niche space, spatial scale and pollinator importance measurement, nonmetric multidimensional scaling (NMDS) was performed on the matrix of interspecific pairwise distances in relative importance of pollinator types. Computations were performed with the function metaMDS in the package vegan (Oksanen et al. 2019) for the R computing environment (R Core Team 2018; all statistical analyses in this paper were carried out using R). In metaMDS the ordination is followed by a rotation via principal components analysis, which consistently ensures that the first axis reflects the principal source of variation in the data and thus enhances the possibility of revealing clusters. Unless otherwise stated the Manhattan distance metric (also known as L1 norm, city block or taxicab metric; Deza and Deza 2013) was used to compute interspecific distances matrices for NMDS and rest of analyses involving distance matrices. The Manhattan metric is better suited than the widely used Euclidean metric for analyses of high dimensionality data as those done here (Hinneburg et al. 2000, Aggarwal et al. 2001, Houle et al. 2010).

Formal statistical analyses of “clusterability”, or whether a meaningful clustered structure can be objectively established in a multidimensional data set (Ackerman and Ben-David 2009, Nowakowska et al. 2015, Simovici and Hua 2019), were conducted at each spatial scale on the set of plant species coordinates in each pollinator niche space (defined by either proportion of visitors or proportion of visits). Dip multimodality tests (Hartigan and Hartigan 1985) were applied to distributions of pairwise distances between species, a method particularly robust to outliers, anomalous topologies and high dimensionality (Adolfsson et al. 2019). These tests evaluated the null hypothesis that the distribution of interspecific distances in pollinator space was unimodal, denoting a single species cluster, against the alternative hypothesis that the distribution had two or more modes, denoting the existence of at least two clusters (Adolfsson et al. 2019; see Brownstein et al. 2019, Simovici and Hua 2019, for examples of applications). The clusterabilitytest function in the clusterability package (Neville and Brownstein 2019) was used for computations, and statistical significance was determined by Monte Carlo simulations with 10^4^ repetitions. Since results of clusterability tests did not support the Fig. 1A scenario, it was important to assess whether the failure to identify species clusters in the pollinator niche spaces considered was an spurious outcome of pooling plant species from different habitats and with different flowering phenologies, which could be partitioning pollinators in time or space. This possibility was examined by simultaneously regressing each matrix of interspecific distance in pollinator composition against corresponding pairwise matrices of distances in blooming time (absolute difference between mean pollinator census dates) and habitat type dissimilarity (simple mismatching coefficient), using the multi.mantel function in the phytools package (Revell 2012). Statistical significance was evaluated using permutations with 10^4^ repetitions.

Simulations were performed to elucidate whether arrangement of plant species within the single cluster found in pollinator niche space matched any of the scenarios depicted in Figs. 1B and 1C. The observed mean pairwise distance between species in each pollinator space was compared with a distribution of means obtained from simulated pollinator composition data. Each simulation consisted of permuting the values in each column of the raw plant species x pollinator type data matrix (number of pollinator individuals or number of flower visits contributed by each pollinator type); computing for each plant species the proportional importance of each pollinator type in the simulated data; obtaining the pairwise distances matrix; and finally computing the mean pairwise distance between species. This simulation procedure created random configurations of proportional pollinator compositions for individual plant species, and thus random arrangements of plants in the corresponding pollinator niche space, but preserved exactly both the observed distributional properties of visitors or visits for every pollinator type, and the absolute and relative numerical importances of the different pollinator types for all plant species combined. A total of 10^4^ simulated species arrangements were computed in each of the four pollinator niche spaces, using proportion of visitors and proportion of visits as measurements of pollinator type importance. To test whether the observed arrangement of plant species in the pollinator niche space departed significantly from the randomly simulated ones, the mean interspecific distance observed was compared with the 0.025 and 0.975 quantiles of the distribution of simulated means.

## Results

The number of orders, families, genera and species of pollinators recorded for each plant species are shown in Appendix S2: Table S1. Most plant species in the sample (*N* = 221) had very taxonomically diverse arrays of insect pollinators. On average (± SE), individual plant species were pollinated by 3.3 ± 0.1 different insect orders (interquartile range, IQR = 3–4), 10.7 ± 0.4 insect families (IQR = 6–14), 16.7 ± 0.8 insect genera (IQR = 7–22), and 16.5 ± 0.8 insect species (IQR = 7–21). This latter figure is an obvious underestimate given the incomplete identification of pollinators to species noted earlier (*Materials and Methods: Data analysis*).

Neither the informal analysis based on nonmetric multidimensional scaling (NMDS) nor the formal one testing clusterability by means of dip tests supported the existence of objectively definable plant species clusters in any of the four pollinator niche spaces considered and regardless of whether analyses referred to the regional or local scales. Irrespective of whether pollinator composition was assessed using proportion of visitors or proportion of visits, the cloud of points in the reduced two-dimensional space obtained with NMDS was roughly centered on the bivariate origin (0, 0), and there was not any indication of clusters or discontinuities suggestive of species groups with contrasting, well-defined pollinator types (Fig. 3). Clusterability analyses using dip tests likewise failed to detect clustered structures underlying the data in any of the pollinator niche spaces or spatial scales considered (Table 2).

**Fig. 3.**
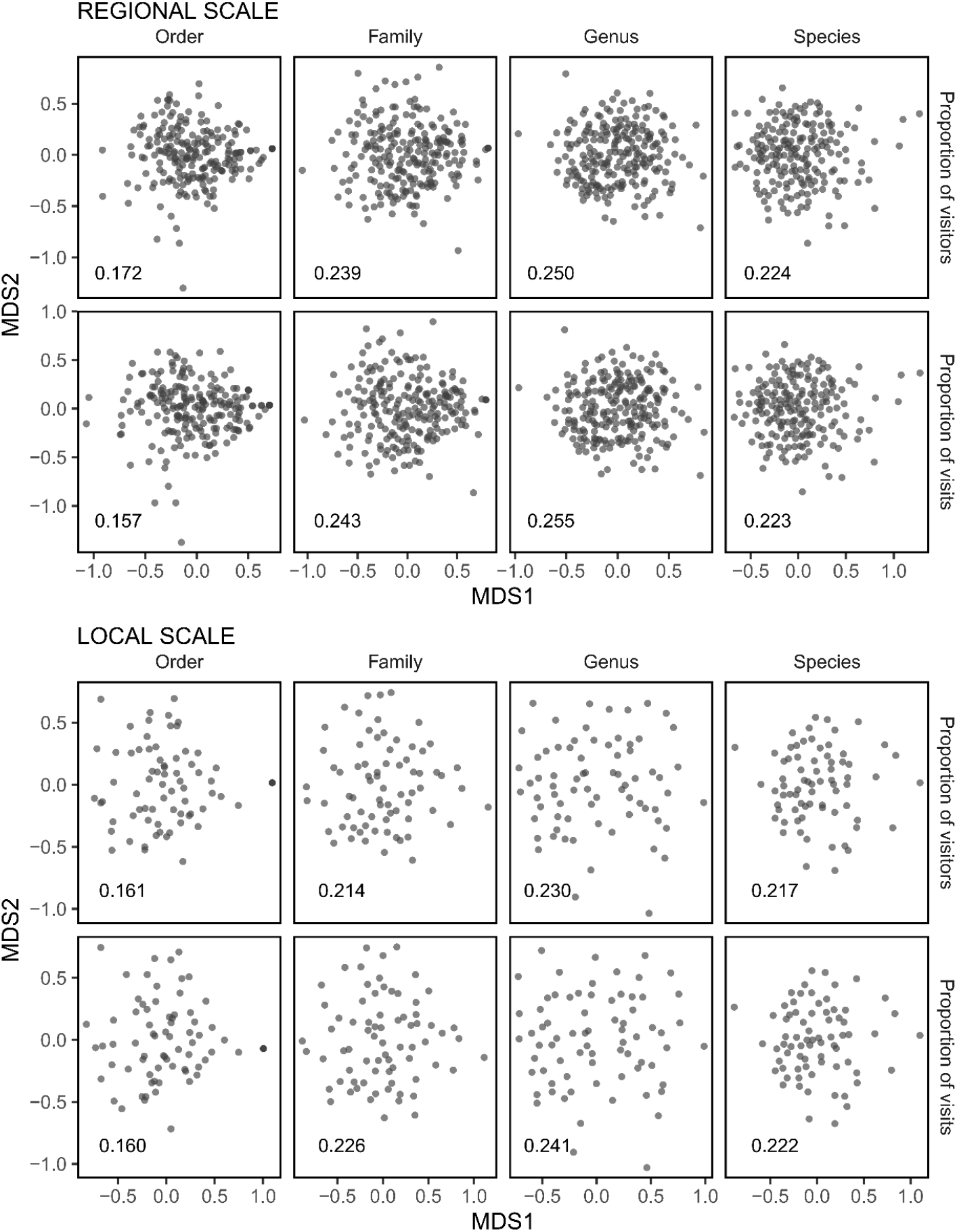
Distribution of plant species on the plane defined by axes from Nonmetric Multidimensional Scaling applied to the matrices of pairwise distances in pollinator composition (MDS1, MDS2). Analyses at the regional scale (two upper rows) consider data from 221 plant species sampled at 45 sites, while those at the local scale (two lower rows) include data from 73 plant species from a single, intensively sampled site (see Fig. 2). Original pairwise distances were computed using two different measurements of relative pollinator importance (proportions of visitors recorded and proportion of flowers visited), and referred to four different pollinator niche spaces whose axes are defined by insect orders, families, genera and species. Stress values, an inverse measurement of goodness of fit, are shown in graphs. Note that meanings of MDS1 and MDS2 axes are not comparable across graphs, since axis pairs were obtained from independent ordination analyses on different data. See Table 1 for sample sizes and dimensionality of the different pollinator niche spaces at the two spatial scales considered.

**Table 2.**
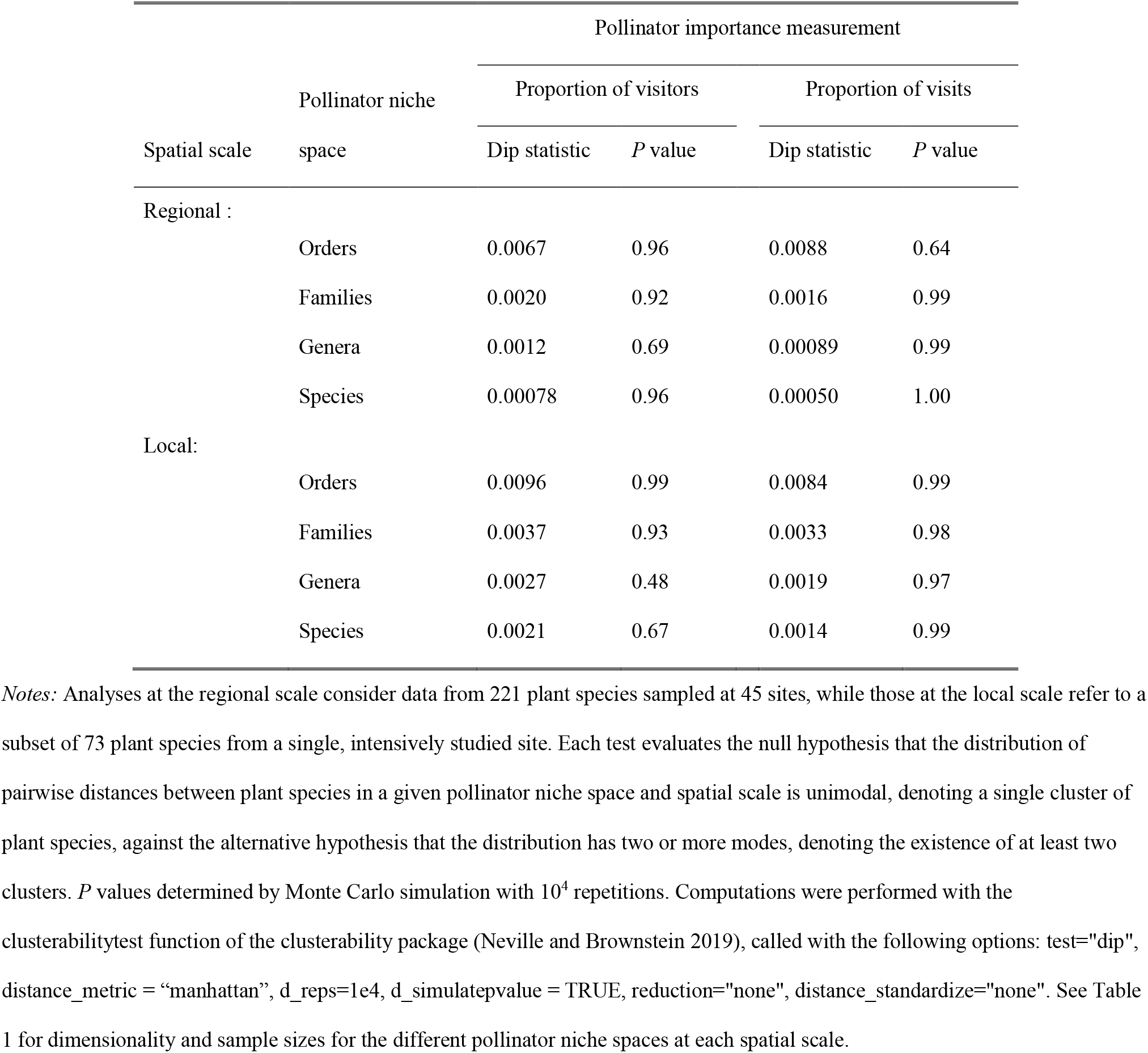
Tests for multimodality of pairwise distances between plant species at two spatial scales and four different pollinator niche spaces whose axes are defined by insect orders, families, genera and species.

|  |  | Pollinator importance measurement |  |  |  |
| --- | --- | --- | --- | --- | --- |
|  | Pollinator niche | Proportion of visitors |  | Proportion of visits |  |
| Spatial scale | space | Dip statistic | *P* value | Dip statistic | *P* value |
| Regional : |  |  |  |  |  |
|  | Orders | 0.0067 | 0.96 | 0.0088 | 0.64 |
|  | Families | 0.0020 | 0.92 | 0.0016 | 0.99 |
|  | Genera | 0.0012 | 0.69 | 0.00089 | 0.99 |
|  | Species | 0.00078 | 0.96 | 0.00050 | 1.00 |
| Local: |  |  |  |  |  |
|  | Orders | 0.0096 | 0.99 | 0.0084 | 0.99 |
|  | Families | 0.0037 | 0.93 | 0.0033 | 0.98 |
|  | Genera | 0.0027 | 0.48 | 0.0019 | 0.97 |
|  | Species | 0.0021 | 0.67 | 0.0014 | 0.99 |

*Notes:* Analyses at the regional scale consider data from 221 plant species sampled at 45 sites, while those at the local scale refer to a subset of 73 plant species from a single, intensively studied site. Each test evaluates the null hypothesis that the distribution of pairwise distances between plant species in a given pollinator niche space and spatial scale is unimodal, denoting a single cluster of plant species, against the alternative hypothesis that the distribution has two or more modes, denoting the existence of at least two clusters. *P* values determined by Monte Carlo simulation with $10^4$ repetitions. Computations were performed with the clusterabilitytest function of the clusterability package (Neville and Brownstein 2019), called with the following options: test="dip", distance\_metric = "manhattan", d\_reps=1e4, d\_simulatepvalue = TRUE, reduction="none", distance\_standardize="none". See Table 1 for dimensionality and sample sizes for the different pollinator niche spaces at each spatial scale.

Multiple Mantel regressions revealed no statistically significant effects of interspecific dissimilarity in flowering time or habitat type on pollinator composition dissimilarity when the axes of pollinator niche space were insect orders, irrespective of whether analyses referred to the regional or local scales (Table 3). When niche spaces were defined by insect families, genera and species, in contrast, statistically significant relationships existed at both the regional and local scales between dissimilarity in pollinator composition and dissimilarity in flowering time (Table 3). Significant relationships between dissimilarity in pollinator composition and habitat dissimilarity occurred only at the regional scale. Regression slopes were positive in all instances of statistically significant relationships, thus denoting that plant species pairs sharing the same habitat type (regional scale) or flowering at similar times of year (regional and local scales) tended to be more similar in pollinator composition at the insect family, genus and species level than those from different habitats or flowering at different times of year (Table 3).

**Table 3.**
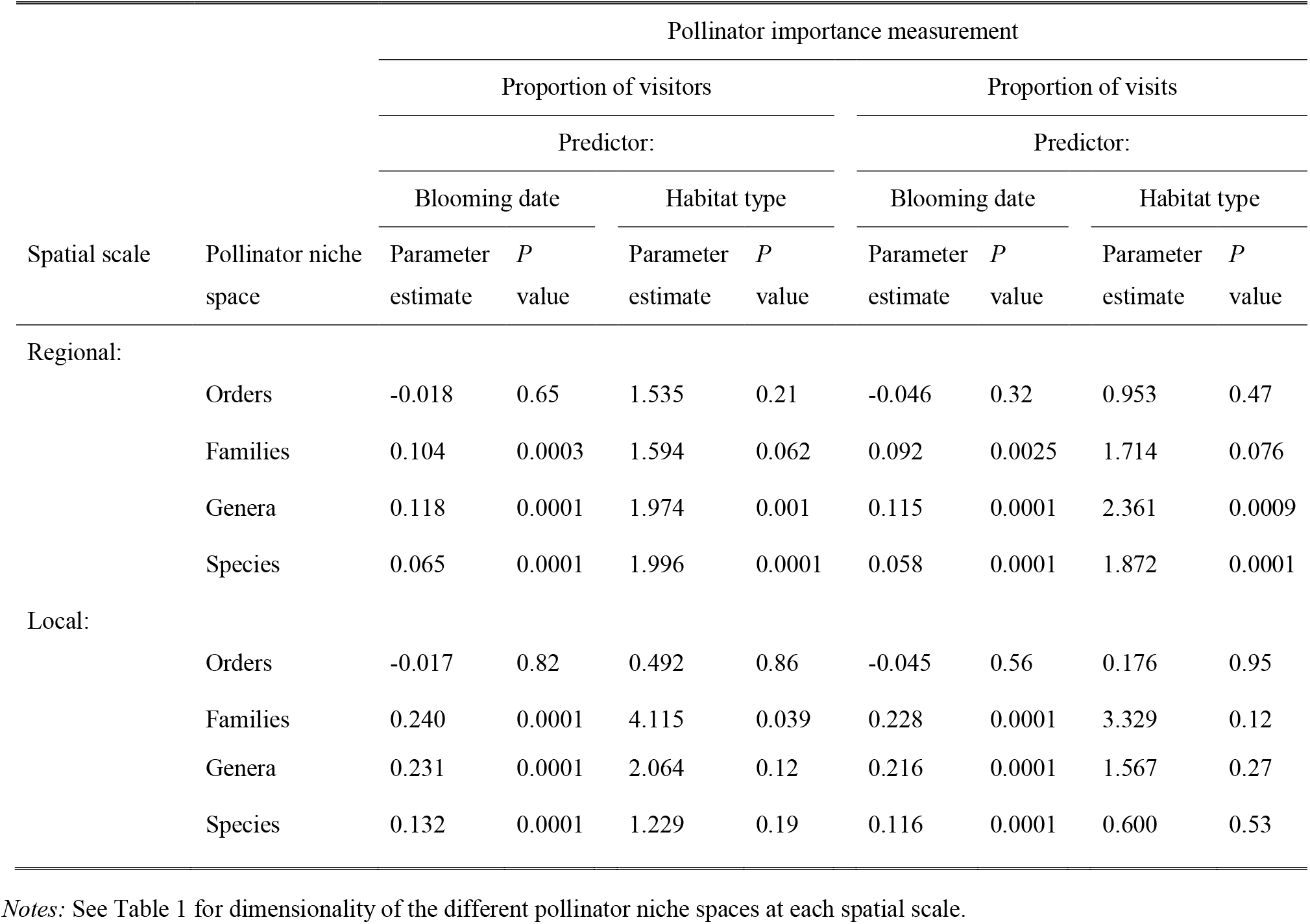
Results of multiple Mantel regressions simultaneously assessing the significance of dissimilarity between plant species in blooming date and habitat type as predictors of distance in pollinator composition, at two spatial scales and using four different pollinator niche spaces whose axes are defined by insect orders, families, genera and species.

Simulations showed that, within the single recognizable cluster, plant species were significantly underdispersed in pollinator niche spaces defined by insect orders, families and genera, and randomly distributed in the niche space defined by insect species. Results were similar for the two spatial scales and the two measurements of pollinator importance (Fig. 4). For order-, family- and genus-defined niche spaces, observed mean of pairwise interspecific distances in pollinator composition was always significantly smaller than simulated values, falling below the lower limit of the range defined by the 0.025–0975 quantiles (Fig. 4). The extent of underdispersion, as intuitively assessed by the disparity between the observed mean and the range of simulated ones, increased from order-through family-to genus-defined pollinator niche spaces.

**Fig. 4.**
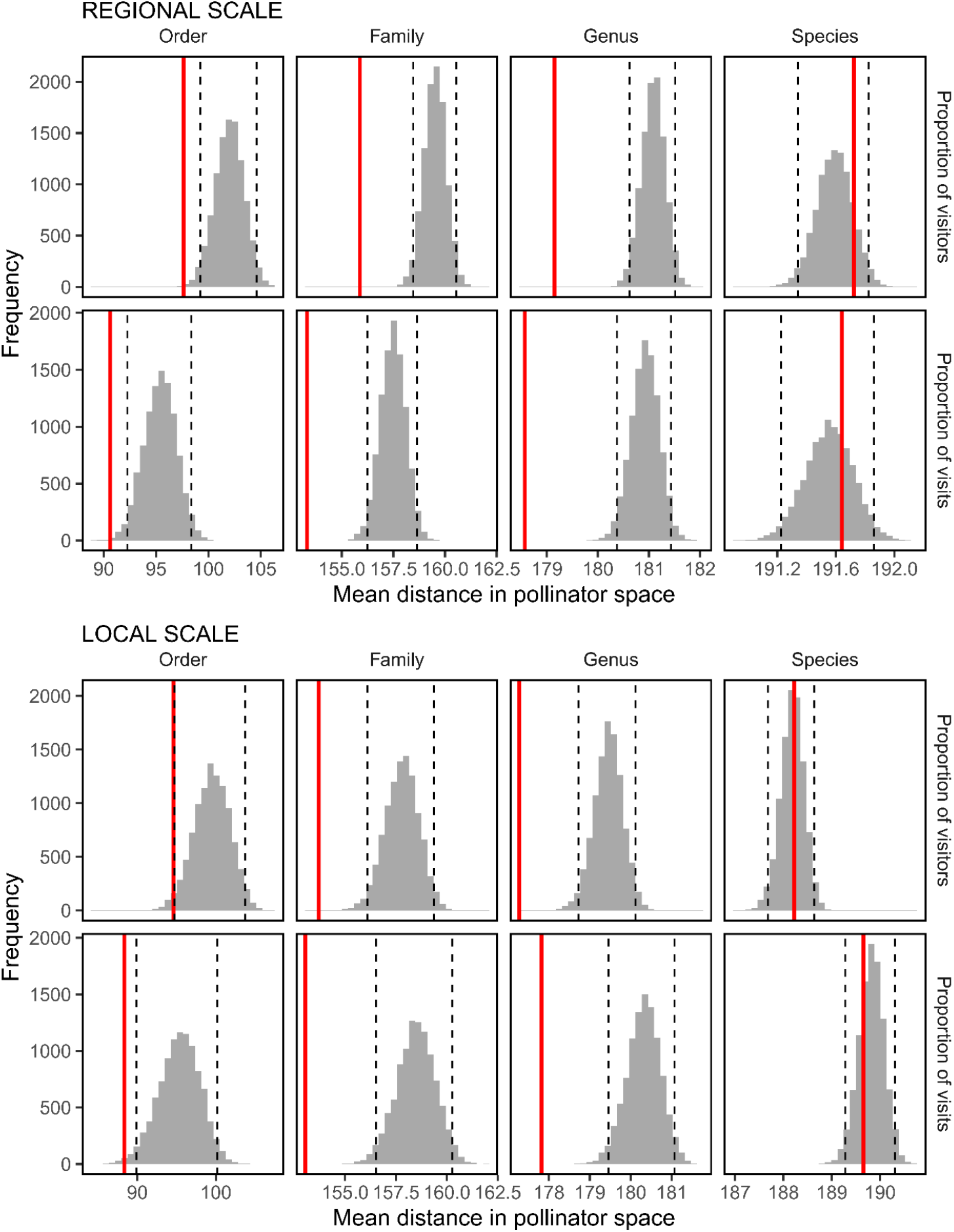
Observed (red lines) and randomly simulated (grey histograms; *N* = 10^4^ simulations) values for the mean pairwise distance between plant species in four different pollinator niche spaces and two spatial scales. Distances were computed for two different measurements of relative pollinator importance (proportions of visitors recorded or proportions of flowers visited). Axes of pollinator niche spaces are defined by insect orders, families, genera and species (see Table 1 for number of dimensions). Analyses at the regional and local scales are based on 221 plant species from 45 sites, and 73 plant species from one site, respectively. The dashed lines denote the 0.025 and 0.975 quantiles of the distributions of simulated mean values.

## Discussion

Considerable effort has been invested to develop reliable methods for grouping sets of objects in multivariate space in such a way that objects falling in the same group are more similar among themselves than to objects in other groups, a technique known as “cluster analysis” (Hennig et al. 2016) widely used in ecological applications. Clustering techniques often focus on the problem of optimally splitting a multivariate data set into *k* ≥ 2 clusters (Halkidi et al. 2016), yet the problem of elucidating whether cohesive and well-differentiated groups of objects actually exist in high dimensionality data, i.e., testing the null hypothesis that *k* = 1 against the alternative *k* ≥ 2, has been rarely investigated due to computational difficulties (the so-called “clusterability problem”; Ackerman and Ben-David 2009, Nowakowska et al. 2015, Simovici and Hua 2019). Following recent technical developments, it has been advised that a prior study of clusterability should become an integral part of any cluster analysis (Adolfsson 2016, Adolfsson et al. 2019, Simovici and Hua 2019). Clusterability tests could be profitably incorporated as a first step in analyses of ecological questions traditionally addressed by means of cluster analysis, so that the existence in multivariate spaces of objectively recognizable clusters of objects (e.g., species, plots, communities) can be rigorously evaluated before proceeding to clustering. This applies also to pollination studies, which have sometimes used clustering to identify plant groupings in multivariate spaces defined by floral traits or pollinator composition (Ollerton et al. 2009, de Jager et al. 2011, Abrahamczyk et al. 2017, Schurr et al. 2019, Vandelook et al. 2019). As exemplified by this study, clusterability analysis allows tests of hypotheses in pollination ecology by providing an objective means of ascertaining whether groups defined by pollinator composition actually exist in a given plant species sample.

The present study failed to identify distinct clusters of plant species in pollinator niche space. Groups with plant species more similar to each other in pollinator composition than to plant species in other groups were not statistically recognizable in the very large species sample of insect-pollinated plants considered. This conclusion emerges as a particularly robust one, since it was consistently attained regardless of the particular taxonomic resolution adopted to define the axes of the pollinator niche space (insect orders, families, genera, species), the spatial scale considered (regional *vs*. local) or the measurement used to quantify pollinator importance (proportion of visitors, proportion of visits). Irrespective of niche space, spatial scale or pollinator importance measurement, variation in pollinator composition across the large plant species sample was thus remarkably continuous. This was largely a consequence of the generalized insect pollination systems exhibited by the vast majority of plant species considered, each of which had pollinators belonging to a substantial number of different insect orders and families. Similar predominance of generalized pollination systems has been reported by other community studies (McCall and Primack 1992, Waser et al. 1996, Bosch et al. 1997), but I am not aware of any previous investigation evaluating the reality of plant species clusters in multidimensional pollinator spaces by means of objective statistical methods. Quantitative studies of plant-pollinator networks have often found “modules” consisting of subsets of plant and animal taxa that are closely linked internally but weakly linked to other modules (Olesen et al. 2007, Dormann and Strauss 2014). Nevertheless, given the loose connection existing between community ecology and network approaches (Blüthgen 2010), the module concept is not strictly comparable to that of species cluster in pollinator niche space envisaged by this study. Plant species clusters associated with “pollination niches” defined on the basis of functional pollinator features (Paw 2013, Phillips et al. 2020) are neither comparable to the neutrally defined clusters in pollination niche space envisaged in this paper.

Pollinators from different insect orders, families, genera or species generally have contrasting morphological and behavioral attributes that can eventually lead to variation in important pollinating features such as, e.g., ability to access floral rewards, daily or seasonal activity rhythm, flower visitation rate, frequency of pollen removal or deposition, or flight distance between flowers. On this basis, taxonomically-defined groupings of insects have been often treated as “functional pollinator groups” related to distinct plant “pollination syndromes”, or suites of floral features that are associated to particular pollinator groups (Fægri and van der Pijl 1979, Fenster et al. 2004, Ollerton et al. 2007, 2009). The relationship between insect taxonomic affiliation and pollination attributes thus lends biological justification for using taxonomic categories to define axes in multidimensional pollinator niche spaces. This study has shown that the arrangement of plant species in various pollinator niche spaces did not match the scenarios postulated on the assumption that interspecific competition is a major force shaping pollinator resource use by plants by favoring pollinator partitioning. Both nonmetric multidimensional scaling and clusterability analyses consistently failed to reveal objectively discernible plant species clusters for any of the levels of pollinator taxonomic resolution, spatial scales or pollinator importance measurements. Furthermore, in the pollinator niche spaces defined by insect orders, families and genera the arrangement of plants was significantly more dense than expected by chance given the observed abundances of the different pollinator types considered. Mantel multiple regressions revealed that plant species sharing the same habitats or blooming at the same time of year were more similar to each other in pollinator composition at the insect family, genus and species levels than those that segregated temporally or spatially, which does not support the possibility that interspecific competition could have favored interspecific divergence between plant species in blooming time of habitat type. The only exceptions to this pattern were the statistically nonsignificant relationships between pollinator composition and habitat dissimilarity at the local scale, which most likely reflects reduced habitat heterogeneity at the limited spatial extent of the single site involved.

When the axes of pollinator niche space were defined by insect species, observed means of pairwise interspecific distances in pollinator composition did not depart significantly from randomly generated means, irrespective of spatial scale or pollinator importance measurement. This result contrasts with the underdispersion found in the rest of niche spaces and possibly reflects random dissimilarities in pollinator composition at the insect species level. This parsimonious interpretation, however, demands caution. Missing species-level identifications in my pollinator sample tended to concentrate in very species-rich genera whose species are difficult to tell apart by external features alone (e.g., *Andrena* and *Lasioglossum* bees; *Phthiria* and *Villa* flies; *Pyrgus* butteflies; *Aplocnemus* beetles). Taxonomically non-random distribution of records with missing species-level identifications could have led to taxonomically biased reduction in niche space dimensionality and distorted plant arrangements in pollinator niche space.

Results of this study suggest that concepts quintessential to competition-laden niche theory, such as limiting similarity, character displacement and competitive exclusion (Chase and Leibold 2003, Johnson and Bronstein 2019), do not seem to apply generally to insect-pollinated plants in my study region in relation to their exploitation of insect pollinator resources. In addition, the absence in my plant species sample of objectively recognizable clusters linked to particular pollinator groups does not support the generality of plant pollination syndromes associated to specialization on particular insect groups (see also Ollerton et al. 2009, 2015). Observed patterns, consisting of plants not showing objectively-definable partitioning of pollinators and being underdispersed in pollinator niche spaces defined at the higher levels of the taxonomic hierarchy, would fall close to the facilitation-dominated extreme in the hypothesized competition-facilitation gradient. In small sets of plant species the competition-facilitation balance in plant-pollinator systems has been shown to vary depending on elements of the ecological scenario such as, e.g., plant species composition, flowering density or pollinator abundance (Moeller 2004, Ghazoul 2006, Ye et al. 2014, Fitch 2017, Eisen et al. 2019). Results of the present study at the plant community level can likewise be interpreted in relation to broad features of plant-pollinator interactions in the study area.

On one side, insect pollinators probably are not a limiting resource there for most insect-pollinated plants, as shown by the high flower visitation rates for most plant species in most habitat types (Herrera 2020), generally weak or inexistent pollen limitation even in plants blooming at the most adverse periods of year (Herrera 1995, 2002, Alonso 2005, Alonso et al. 2012, 2013), and increased pollinator abundance in recent times in parallel to the long-term warming trend exhibited by the regional climate (Herrera 2019). Since abundance of pollinators in the study region is most likely related to the preservation of its natural habitats, it seems plausible to postulate that anthropogenic disturbances of natural habitats should favor the presence of plant species clusters in pollinator niche space. And on the other side, sharing of individual pollinators by different plant species seems a widespread phenomenon, as denoted by the frequent occurrence of heterospecific pollen grains in the stigmas of most species (Arceo-Gómez et al. 2018), common field observations of insect individuals interspersing visits to flowers of different species in the same foraging bout, and close-up photographs often revealing conspicuous mixed pollen loads on pollinator bodies (C. M. Herrera, *unpublished data*). Taken together, all these observation are consistent with the interpretation that the unclusterable, predominantly underdispersed arrangement of plant species in pollinator niche space found in this study reflects a prevailing role of interspecific facilitation as the organizing driver of plant-pollinator systems in the undisturbed montane habitats studied. Future investigations conducted in plant communities from other latitudes and continents, particularly where plants exploit taxonomically broader pollinator assemblages (e.g., including diurnal and nocturnal vertebrates), should help to evaluate the generality of the results of this study, revealing whether plant species are arranged differently in disturbed habitats or in pollinator niche spaces whose axes are defined by a broader variety of animal groups (e.g., insects, birds and mammals).

## Concluding Remarks

Excepting a few classical monographs considering complete or nearly complete regional plant–pollinator assemblages (Müller 1883, Robertson 1928, Moldenke 1976), the vast majority of recent pollination community studies have examined only small numbers of plant species at a time (see, e.g., Ollerton 2017:Appendix 2, and datasets included in the bipartite package for R, Dormann et al. 2008). Small species sets drawn from much larger pools of insect-pollinated species provide suitable systems for investigating whether a particular phenomenon occurs or not (e.g., testing for pollinator-mediated competition or facilitation in a well-chosen group of closely-related species; see references in *Introduction*), but are intrisically unable to address more ambitious, broadly-encompassing questions bearing on the strength or frequency with which the phenomenon of interest operates in a particular ecological context. In addition, unacknowledged biases in the selection of pollination study systems can lead to small plant species samples rarely, if ever, representing truly scaled-down, taxonomically- and ecologically-unbiased versions of whole plant communities, possibly leading to distorted representations of natural patterns (Ollerton et al. 2015). Place-based ecological research pursuing “general understanding through the sort of detailed understanding of a particular place” (Price and Billick 2010, p. 4) has been only infrequently adopted by pollination studies. As exemplified by the present investigation (see also Herrera 2020), comprehensive studies of complete or nearly complete plant communities possess an unexplored potential for testing novel hypotheses (or old hypotheses in novel ways) and contributing realistic insights to the understanding of the ecology and evolution of plant-pollinator systems.

## Acknowledgments

I am deeply indebted to all insect taxonomists that helped over the years with insect identifications, a complete list of which was presented in Herrera (2020: *Acknowledgments*). Consejería de Medio Ambiente, Junta de Andalucía, granted permission to work in the Sierra de Cazorla and provided invaluable facilities there. Randall J. Mitchell, James D. Thomson and two anonymous reviewers contributed useful comments that improved considerably the manuscript. Cogent criticisms by Mónica Medrano at an early stage of analyses helped to refine my thinking and clarify the analytical approach. The research reported in this paper received no specific grant from any funding agency.

## Supporting Information

### Appendix S1

Plant species studied, habitat types, flowering time and pollinator sampling effort.

### Appendix S2

Number of taxonomic categories of insect pollinators recorded per plant species.

