## Appendix S1 for "Unclusterable, underdispersed arrangement of insect-pollinated plants in pollinator niche space"

**Supporting Information.** Carlos M. Herrera. Unclusterable, underdispersed arrangement of insect-pollinated plants in pollinator niche space. *Ecology*.

**Appendix S1 Plant species studied, habitat types, flowering time and pollinator sampling effort.**

TABLE S1. Plant species ($N$ = 221) sampled for pollinator visitation during 1997-2019 in the Sierra de Cazorla mountains, southeastern Spain, and data on habitat type, flowering phenology and pollinator sampling effort.

| Species [1] | Family | Habitat type [2] | Flowering time [3] | Pollinator censuses | Sampling dates | Pollinator individuals | Flower visits |
|---|---|---|---|---|---|---|---|
| *Achillea odorata* * | Asteraceae | For-Cl | 175 | 147 | 8 | 588 | 757 |
| *Acinos alpinus* | Lamiaceae | For-Cl | 154 | 94 | 2 | 73 | 457 |
| *Allium roseum* | Alliaceae | Gra | 140 | 205 | 4 | 153 | 456 |
| *Allium scorodoprasum* * | Alliaceae | Gra | 212 | 215 | 7 | 526 | 1343 |
| *Allium sphaerocephalon* * | Alliaceae | For-Cl | 196 | 85 | 4 | 227 | 345 |
| *Alyssum simplex* | Brassicaceae | Gra | 108 | 85 | 3 | 22 | 143 |
| *Amelanchier ovalis* * | Rosaceae | For-Cl | 145 | 75 | 4 | 165 | 221 |
| *Anagallis monelli* * | Primulaceae | Dms | 170 | 100 | 5 | 70 | 221 |
| *Anarrhinum laxiflorum* | Veronicaceae | Scle | 175 | 50 | 1 | 42 | 520 |
| *Andryala ragusina* * | Asteraceae | Dis | 201 | 70 | 3 | 140 | 343 |
| *Anthemis pedunculata* | Asteraceae | Gra | 153 | 95 | 4 | 93 | 123 |

| | | | | | | | |
|---|---|---|---|---|---|---|---|
| *Anthericum liliago* | Asphodelaceae | Dms | 171 | 76 | 2 | 136 | 253 |
| *Anthyllis vulneraria* | Fabaceae | For-In | 149 | 140 | 4 | 93 | 751 |
| *Antirrhinum australe* * | Veronicaceae | Cli | 172 | 80 | 4 | 8 | 14 |
| *Aphyllanthes monspeliensis* * | Aphyllanthaceae | For-Cl | 157 | 200 | 6 | 218 | 677 |
| *Aquilegia cazorlensis* | Ranunculaceae | Cli | 163 | 354 | 10 | 151 | 896 |
| *Aquilegia vulgaris* | Ranunculaceae | Str | 151 | 298 | 8 | 155 | 547 |
| *Arbutus unedo* | Ericaceae | Scle | 364 | 110 | 2 | 44 | 159 |
| *Arenaria armerina* * | Caryophyllaceae | Dol | 195 | 96 | 4 | 66 | 320 |
| *Arenaria modesta* | Caryophyllaceae | Dol | 162 | 110 | 4 | 28 | 93 |
| *Arenaria obtusiflora* * | Caryophyllaceae | Dol | 169 | 100 | 3 | 60 | 172 |
| *Armeria filicaulis* * | Plumbaginaceae | Dms | 153 | 161 | 7 | 191 | 260 |
| *Armeria villosa* | Plumbaginaceae | Gra | 170 | 75 | 2 | 132 | 188 |
| *Asphodelus cerasiferus* | Asphodelaceae | For-Cl | 145 | 136 | 2 | 284 | 1165 |
| *Astragalus bourgaeanus* * | Fabaceae | Dms | 133 | 85 | 5 | 8 | 43 |
| *Astragalus incanus* | Fabaceae | Gra | 131 | 78 | 4 | 46 | 259 |
| *Atropa baetica* | Solanaceae | For-Cl | 182 | 70 | 2 | 115 | 382 |
| *Bellis perennis* | Asteraceae | Gra | 137 | 110 | 2 | 63 | 97 |
| *Bellis sylvestris* | Asteraceae | Gra | 121 | 100 | 3 | 87 | 156 |
| *Berberis hispanica* | Berberidaceae | For-Cl | 159 | 175 | 4 | 288 | 975 |
| *Biscutella laxa* | Brassicaceae | Dol | 124 | 120 | 2 | 81 | 130 |
| *Calamintha nepeta* | Lamiaceae | For-In | 233 | 80 | 3 | 50 | 231 |

| | | | | | | | |
|---|---|---|---|---|---|---|---|
| *Campanula dieckii* * | Campanulaceae | For-Cl | 176 | 130 | 5 | 89 | 289 |
| *Campanula mollis* * | Campanulaceae | Cli | 205 | 110 | 4 | 37 | 54 |
| *Carduncellus monspelliensium* | Asteraceae | Dms | 179 | 115 | 2 | 123 | 259 |
| *Carduus platypus* | Asteraceae | Dis | 156 | 65 | 2 | 167 | 305 |
| *Carduus tenuiflorus* | Asteraceae | Dis | 168 | 81 | 3 | 114 | 275 |
| *Carlina hispanica* | Asteraceae | Dis | 228 | 145 | 6 | 268 | 464 |
| *Carlina racemosa* | Asteraceae | Dis | 238 | 140 | 4 | 286 | 825 |
| *Catananche caerulea* | Asteraceae | For-In | 172 | 150 | 6 | 433 | 655 |
| *Centaurea calcitrapa* | Asteraceae | Dis | 189 | 126 | 4 | 630 | 1137 |
| *Centaurea graminifolia* * | Asteraceae | Dms | 177 | 105 | 3 | 132 | 257 |
| *Cerastium gibraltaricum* * | Caryophyllaceae | Gra | 163 | 85 | 3 | 42 | 137 |
| *Chaenorhinum macropodum* | Veronicaceae | Dol | 155 | 85 | 4 | 10 | 78 |
| *Chiliadenus glutinosus* * | Asteraceae | Dol | 216 | 83 | 4 | 284 | 613 |
| *Chondrilla juncea* * | Asteraceae | Dis | 228 | 175 | 7 | 460 | 1119 |
| *Cirsium acaule* * | Asteraceae | Gra | 225 | 105 | 3 | 139 | 175 |
| *Cirsium monspessulanum* | Asteraceae | Str | 222 | 75 | 4 | 274 | 591 |
| *Cirsium pyrenaicum* | Asteraceae | Str | 224 | 140 | 6 | 365 | 684 |
| *Cirsium vulgare* | Asteraceae | Dis | 227 | 110 | 2 | 167 | 332 |
| *Cistus albidus* | Cistaceae | Scle | 113 | 100 | 3 | 317 | 540 |
| *Cistus monspeliensis* | Cistaceae | Scle | 152 | 154 | 4 | 1040 | 1370 |
| *Cistus salviifolius* | Cistaceae | Scle | 114 | 81 | 3 | 324 | 431 |

| | | | | | | | |
|---|---|---|---|---|---|---|---|
| *Cleonia lusitanica* | Lamiaceae | Scle | 169 | 60 | 2 | 77 | 358 |
| *Clinopodium vulgare* * | Lamiaceae | For-In | 191 | 70 | 3 | 57 | 176 |
| *Conopodium arvense* | Apiaceae | For-In | 160 | 110 | 4 | 70 | 202 |
| *Convolvulus arvensis* * | Convolvulaceae | Dis | 193 | 80 | 3 | 184 | 273 |
| *Coris monspeliensis* | Primulaceae | Dms | 166 | 80 | 3 | 47 | 292 |
| *Crataegus monogyna* | Rosaceae | For-Cl | 175 | 90 | 3 | 121 | 418 |
| *Crepis albida* | Asteraceae | Dol | 158 | 67 | 3 | 121 | 170 |
| *Crocus nevadensis* | Iridaceae | Gra | 39 | 90 | 2 | 35 | 41 |
| *Crocus nudiflorus* * | Iridaceae | Gra | 274 | 100 | 3 | 73 | 120 |
| *Cuscuta triumvirati* * | Convolvulaceae | Dms | 199 | 105 | 4 | 157 | 475 |
| *Daphne gnidium* | Thymelaeaceae | Scle | 227 | 100 | 3 | 38 | 155 |
| *Daphne laureola* | Thymelaeaceae | For-In | 93 | 107 | 3 | 85 | 177 |
| *Digitalis obscura* | Veronicaceae | Scle | 161 | 180 | 6 | 398 | 1561 |
| *Draba hispanica* * | Brassicaceae | Cli | 115 | 102 | 2 | 50 | 127 |
| *Echinospartum boissieri* * | Fabaceae | Dms | 173 | 95 | 4 | 37 | 126 |
| *Echium flavum* | Boraginaceae | Dis | 146 | 160 | 4 | 134 | 1243 |
| *Erinacea anthyllis* | Fabaceae | Dms | 145 | 136 | 2 | 95 | 761 |
| *Erodium cazorlanum* | Geraniaceae | Dol | 158 | 75 | 3 | 12 | 20 |
| *Erodium cheilanthifolium* | Geraniaceae | Dms | 158 | 105 | 3 | 55 | 103 |
| *Erodium cicutarium* | Geraniaceae | Dis | 113 | 71 | 3 | 63 | 208 |
| *Erophila verna* | Brassicaceae | Gra | 94 | 90 | 3 | 28 | 184 |

| | | | | | | | |
|---|---|---|---|---|---|---|---|
| *Eryngium campestre* | Apiaceae | Dis | 223 | 250 | 9 | 746 | 1620 |
| *Eryngium dilatatum* * | Apiaceae | For-Cl | 198 | 200 | 7 | 443 | 1070 |
| *Erysimum cazorlense* | Brassicaceae | Dol | 148 | 125 | 4 | 28 | 95 |
| *Erysimum medio-hispanicum* | Brassicaceae | For-Cl | 153 | 100 | 3 | 71 | 292 |
| *Euphorbia nicaeensis* | Euphorbiaceae | For-Cl | 174 | 140 | 4 | 406 | 863 |
| *Filipendula vulgaris* * | Rosaceae | Gra | 171 | 72 | 3 | 117 | 145 |
| *Fumana baetica* * | Cistaceae | Dol | 169 | 160 | 8 | 166 | 351 |
| *Fumana paradoxa* | Cistaceae | Dol | 182 | 106 | 2 | 59 | 155 |
| *Fumana procumbens* | Cistaceae | Dol | 175 | 101 | 3 | 22 | 51 |
| *Gagea soleirolii* * | Liliaceae | For-Cl | 116 | 108 | 2 | 115 | 245 |
| *Galatella linosyris* * | Asteraceae | For-Cl | 262 | 120 | 4 | 116 | 329 |
| *Galium verum* * | Rubiaceae | Gra | 194 | 85 | 4 | 124 | 587 |
| *Genista longipes* | Fabaceae | Dol | 160 | 112 | 3 | 19 | 92 |
| *Genista pseudopilosa* * | Fabaceae | Dms | 172 | 100 | 3 | 15 | 119 |
| *Genista scorpius* | Fabaceae | Dis | 103 | 112 | 3 | 51 | 166 |
| *Geranium molle* | Geraniaceae | Dis | 150 | 102 | 3 | 55 | 162 |
| *Geum sylvaticum* | Rosaceae | For-In | 123 | 122 | 3 | 63 | 89 |
| *Gladiolus illyricus* * | Iridaceae | Gra | 175 | 190 | 7 | 269 | 1157 |
| *Globularia spinosa* | Globulariaceae | Dol | 141 | 110 | 3 | 25 | 40 |
| *Gymnadenia conopsea* | Orchidaceae | Str | 160 | 110 | 3 | 14 | 227 |
| *Hedera helix* * | Araliaceae | Cli | 276 | 153 | 4 | 245 | 889 |

| | | | | | | | |
|---|---|---|---|---|---|---|---|
| *Helianthemum apenninum* | Cistaceae | Dms | 150 | 165 | 5 | 260 | 374 |
| *Helianthemum cinereum* | Cistaceae | Dol | 153 | 100 | 4 | 51 | 340 |
| *Helianthemum oelandicum* * | Cistaceae | Dms | 145 | 167 | 6 | 214 | 349 |
| *Helleborus foetidus* | Ranunculaceae | For-In | 63 | 1786 | 21 | 241 | 698 |
| *Hepatica nobilis* | Ranunculaceae | For-In | 109 | 90 | 2 | 11 | 34 |
| *Himantoglossum hircinum* * | Orchidaceae | For-Cl | 147 | 66 | 4 | 11 | 26 |
| *Hormathophylla baetica* | Brassicaceae | Dol | 112 | 41 | 2 | 37 | 123 |
| *Hormathophylla spinosa* | Brassicaceae | Dms | 176 | 90 | 3 | 95 | 509 |
| *Hyacinthoides reverchonii* * | Hyacinthaceae | Cli | 124 | 105 | 4 | 24 | 67 |
| *Hypericum caprifolium* | Clusiaceae | Str | 198 | 76 | 3 | 69 | 673 |
| *Hypericum ericoides* | Clusiaceae | Cli | 199 | 80 | 3 | 31 | 34 |
| *Hypericum perforatum* * | Clusiaceae | For-Cl | 193 | 140 | 9 | 309 | 1071 |
| *Iberis carnosa* | Brassicaceae | Dol | 106 | 125 | 3 | 19 | 94 |
| *Inula montana* * | Asteraceae | Dms | 170 | 155 | 6 | 519 | 929 |
| *Iris foetidissima* | Iridaceae | Str | 170 | 90 | 3 | 30 | 201 |
| *Iris planifolia* | Iridaceae | Gra | 68 | 100 | 2 | 10 | 22 |
| *Jasonia tuberosa* * | Asteraceae | Str | 218 | 70 | 3 | 262 | 479 |
| *Jonopsidium prolongoi* | Brassicaceae | Gra | 121 | 97 | 3 | 55 | 192 |
| *Jurinea humilis* | Asteraceae | Dms | 166 | 92 | 3 | 53 | 81 |
| *Klasea nudicaulis* * | Asteraceae | Gra | 171 | 75 | 3 | 115 | 205 |
| *Klasea pinnatifida* * | Asteraceae | For-Cl | 169 | 165 | 5 | 361 | 635 |

| | | | | | | | |
|---|---|---|---|---|---|---|---|
| *Knautia subscaposa* | Dipsacaceae | Gra | 167 | 177 | 4 | 376 | 680 |
| *Lamium amplexicaule* | Lamiaceae | Dis | 112 | 76 | 2 | 42 | 310 |
| *Lavandula latifolia* | Lamiaceae | Scle | 220 | 818 | 38 | 1012 | 6185 |
| *Leontodon longirrostris* | Asteraceae | Gra | 174 | 65 | 3 | 269 | 456 |
| *Leucanthemopsis pallida* | Asteraceae | Dol | 132 | 70 | 4 | 88 | 162 |
| *Linaria aeruginea* | Veronicaceae | Dol | 155 | 71 | 4 | 7 | 13 |
| *Linaria verticillata* * | Veronicaceae | Cli | 163 | 75 | 3 | 7 | 31 |
| *Linaria viscosa* | Veronicaceae | Dol | 154 | 105 | 3 | 50 | 531 |
| *Linum bienne* * | Linaceae | Gra | 177 | 172 | 7 | 127 | 656 |
| *Linum narbonense* * | Linaceae | For-Cl | 158 | 85 | 5 | 24 | 47 |
| *Linum tenue* | Linaceae | Gra | 171 | 190 | 6 | 350 | 680 |
| *Lithodora fruticosa* * | Boraginaceae | Dms | 154 | 185 | 7 | 113 | 895 |
| *Lonicera arborea* | Caprifoliaceae | For-Cl | 173 | 65 | 2 | 106 | 503 |
| *Lysimachia ephemerum* | Primulaceae | Str | 183 | 200 | 5 | 444 | 1677 |
| *Lythrum junceum* | Lythraceae | Str | 166 | 97 | 4 | 83 | 222 |
| *Lythrum salicaria* | Lythraceae | Str | 238 | 110 | 3 | 88 | 385 |
| *Mantisalca salmantica* * | Asteraceae | For-Cl | 199 | 160 | 6 | 255 | 508 |
| *Marrubium supinum* | Lamiaceae | Dms | 164 | 160 | 7 | 280 | 3427 |
| *Mentha aquatica* | Lamiaceae | Str | 228 | 110 | 3 | 123 | 223 |
| *Merendera montana* * | Colchicaceae | Gra | 248 | 120 | 4 | 88 | 143 |
| *Narcissus bujei* | Amaryllidaceae | Scle | 65 | 205 | 4 | 112 | 191 |

| | | | | | | | |
|---|---|---|---|---|---|---|---|
| *Narcissus cuatrecasasii* | Amaryllidaceae | For-In | 112 | 214 | 5 | 56 | 145 |
| *Narcissus hedraeanthus* | Amaryllidaceae | Gra | 86 | 205 | 7 | 120 | 371 |
| *Narcissus longispathus* | Amaryllidaceae | Str | 90 | 295 | 8 | 288 | 415 |
| *Narcissus triandrus* | Amaryllidaceae | For-In | 113 | 125 | 3 | 15 | 50 |
| *Nepeta tuberosa* | Lamiaceae | Str | 172 | 60 | 2 | 80 | 704 |
| *Ononis pusilla* | Fabaceae | Dol | 158 | 79 | 3 | 32 | 107 |
| *Ononis spinosa* | Fabaceae | Gra | 204 | 100 | 3 | 41 | 212 |
| *Orchis coriophora* * | Orchidaceae | Gra | 175 | 120 | 2 | 97 | 475 |
| *Origanum virens* | Lamiaceae | For-Cl | 198 | 77 | 3 | 77 | 365 |
| *Ornithogalum umbellatum* | Hyacinthaceae | For-Cl | 142 | 160 | 6 | 164 | 359 |
| *Orobanche haenseleri* | Orobanchaceae | For-In | 180 | 96 | 3 | 55 | 162 |
| *Papaver dubium* * | Papaveraceae | Dis | 177 | 85 | 3 | 105 | 199 |
| *Parnassia palustris* | Saxifragaceae | Str | 263 | 70 | 2 | 35 | 39 |
| *Petrorhagia nanteuilii* * | Caryophyllaceae | For-Cl | 178 | 70 | 4 | 123 | 305 |
| *Phlomis herba-venti* | Lamiaceae | Gra | 183 | 150 | 4 | 294 | 1028 |
| *Phlomis lychnitis* | Lamiaceae | Scle | 177 | 151 | 2 | 124 | 705 |
| *Picnomon acarna* | Asteraceae | Dis | 233 | 125 | 4 | 229 | 406 |
| *Pilosella pseudopilosella* * | Asteraceae | Gra | 175 | 66 | 4 | 222 | 405 |
| *Pistorinia hispanica* | Crassulaceae | Dol | 170 | 161 | 5 | 86 | 1157 |
| *Plumbago europaea* * | Plumbaginaceae | Dis | 226 | 205 | 8 | 407 | 2270 |
| *Polygala boissieri* | Polygalaceae | For-In | 146 | 100 | 5 | 19 | 75 |

| | | | | | | | |
|---|---|---|---|---|---|---|---|
| *Polygonatum odoratum* | Ruscaceae | For-In | 142 | 138 | 4 | 13 | 84 |
| *Potentilla caulescens* | Rosaceae | Cli | 225 | 125 | 3 | 122 | 573 |
| *Potentilla reptans* * | Rosaceae | Gra | 132 | 100 | 4 | 89 | 187 |
| *Primula acaulis* | Primulaceae | For-In | 111 | 153 | 5 | 27 | 89 |
| *Prolongoa hispanica* | Asteraceae | Dol | 165 | 88 | 3 | 62 | 82 |
| *Prunus mahaleb* * | Rosaceae | For-Cl | 137 | 120 | 4 | 149 | 463 |
| *Pterocephalus spathulatus* | Dipsacaceae | Dol | 193 | 120 | 3 | 154 | 310 |
| *Ptilostemon hispanicus* | Asteraceae | Dis | 224 | 65 | 3 | 159 | 326 |
| *Ranunculus bulbosus* | Ranunculaceae | Str | 155 | 100 | 3 | 183 | 257 |
| *Ranunculus ficaria* * | Ranunculaceae | For-In | 127 | 108 | 3 | 57 | 64 |
| *Ranunculus malessanus* | Ranunculaceae | Gra | 120 | 95 | 3 | 112 | 154 |
| *Ranunculus paludosus* * | Ranunculaceae | Gra | 131 | 105 | 4 | 136 | 198 |
| *Ranunculus repens* | Ranunculaceae | Gra | 167 | 100 | 3 | 334 | 630 |
| *Rhagadiolus edulis* | Asteraceae | For-In | 165 | 95 | 3 | 49 | 126 |
| *Roemeria argemone* | Papaveraceae | Gra | 144 | 90 | 4 | 45 | 63 |
| *Rosa canina* | Rosaceae | For-Cl | 160 | 75 | 3 | 174 | 334 |
| *Rosa micrantha* * | Rosaceae | For-Cl | 181 | 145 | 7 | 421 | 648 |
| *Rosa sicula* * | Rosaceae | For-Cl | 175 | 80 | 3 | 65 | 122 |
| *Rosmarinus officinalis* | Lamiaceae | Scle | 140 | 166 | 4 | 239 | 2138 |
| *Rubus ulmifolius* | Rosaceae | For-Cl | 190 | 95 | 2 | 524 | 962 |
| *Salvia lavandulifolia* | Lamiaceae | Scle | 185 | 72 | 2 | 77 | 549 |

| | | | | | | | |
|---|---|---|---|---|---|---|---|
| *Salvia verbenaca* | Lamiaceae | Dis | 133 | 90 | 3 | 52 | 159 |
| *Santolina rosmarinifolia* * | Asteraceae | Dms | 192 | 130 | 8 | 427 | 1025 |
| *Saponaria ocymoides* | Caryophyllaceae | For-In | 153 | 141 | 2 | 109 | 1170 |
| *Satureja intricata* * | Lamiaceae | Dms | 193 | 147 | 6 | 220 | 1148 |
| *Saxifraga haenseleri* | Saxifragaceae | Dol | 134 | 90 | 4 | 77 | 120 |
| *Saxifraga carpetana* * | Saxifragaceae | Gra | 132 | 77 | 4 | 37 | 334 |
| *Scabiosa andryaefolia* | Dipsacaceae | For-Cl | 182 | 62 | 2 | 280 | 492 |
| *Scilla paui* | Hyacinthaceae | Gra | 121 | 90 | 3 | 56 | 207 |
| *Scorzonera albicans* * | Asteraceae | Dol | 160 | 70 | 5 | 13 | 24 |
| *Sedum album* * | Crassulaceae | Dms | 185 | 165 | 5 | 595 | 952 |
| *Sedum dasyphyllum* | Crassulaceae | Cli | 193 | 110 | 3 | 76 | 243 |
| *Sedum mucizonia* | Crassulaceae | Dol | 172 | 100 | 3 | 47 | 283 |
| *Sedum sediforme* * | Crassulaceae | Dms | 200 | 95 | 3 | 208 | 714 |
| *Senecio doria* | Asteraceae | Str | 222 | 74 | 4 | 116 | 354 |
| *Senecio malacitanus* | Asteraceae | Dis | 299 | 115 | 3 | 174 | 436 |
| *Seseli montanum* * | Apiaceae | For-Cl | 266 | 120 | 4 | 87 | 305 |
| *Sideritis incana* | Lamiaceae | Dol | 178 | 130 | 6 | 155 | 2035 |
| *Silene colorata* | Caryophyllaceae | Dis | 155 | 92 | 4 | 10 | 37 |
| *Silene psammitis* | Caryophyllaceae | Dol | 149 | 132 | 4 | 111 | 409 |
| *Sisymbrella aspera* * | Brassicaceae | Str | 171 | 180 | 8 | 133 | 408 |
| *Sisymbrium crassifolium* | Brassicaceae | For-Cl | 146 | 70 | 3 | 46 | 182 |

| | | | | | | | |
|---|---|---|---|---|---|---|---|
| *Solidago virgaurea* | Asteraceae | For-In | 235 | 100 | 3 | 42 | 101 |
| *Sonchus aquatilis* | Asteraceae | Str | 232 | 91 | 3 | 283 | 577 |
| *Stachys officinalis* * | Lamiaceae | Gra | 190 | 138 | 8 | 334 | 1799 |
| *Taraxacum laevigatum* | Asteraceae | Gra | 145 | 120 | 3 | 144 | 180 |
| *Teucrium aureum* * | Lamiaceae | Dms | 180 | 142 | 4 | 183 | 1801 |
| *Teucrium rotundifolium* * | Lamiaceae | Cli | 175 | 181 | 6 | 69 | 937 |
| *Teucrium webbianum* | Lamiaceae | Dms | 192 | 81 | 4 | 84 | 472 |
| *Thymus mastichina* | Lamiaceae | Scle | 168 | 160 | 5 | 199 | 1574 |
| *Thymus orospedanus* | Lamiaceae | Dms | 153 | 163 | 4 | 72 | 883 |
| *Thymus serpylloides* | Lamiaceae | Dms | 168 | 75 | 3 | 73 | 952 |
| *Trifolium campestre* * | Fabaceae | Gra | 179 | 82 | 3 | 70 | 463 |
| *Ulex parviflorus* | Fabaceae | Scle | 88 | 125 | 3 | 147 | 578 |
| *Urginea maritima* | Hyacinthaceae | Scle | 236 | 220 | 10 | 575 | 1400 |
| *Valeriana tuberosa* | Valerianaceae | Gra | 136 | 165 | 6 | 81 | 187 |
| *Verbascum giganteum* | Scrophulariaceae | Dis | 218 | 115 | 3 | 139 | 526 |
| *Verbascum sinuatum* | Scrophulariaceae | Dis | 217 | 115 | 3 | 128 | 370 |
| *Viburnum tinus* | Adoxaceae | Scle | 111 | 90 | 3 | 115 | 235 |
| *Vicia onobrychioides* * | Fabaceae | For-Cl | 162 | 65 | 4 | 79 | 473 |
| *Vicia pseudocracca* | Fabaceae | For-Cl | 154 | 73 | 2 | 61 | 347 |
| *Viola cazorlensis* | Violaceae | Cli | 151 | 123 | 6 | 6 | 38 |
| *Viola odorata* | Violaceae | For-In | 112 | 198 | 5 | 31 | 101 |

[1] Asterisks denote species sampled in the intensively-studied Nava de las Correhuelas site ($N$ = 73), included in the local scale analyses.

[2] Mean date of pollinator censuses, expressed as days from 1 January.

[3] Habitat types: Cli, Cliffs; Dis, Disturbances; Dol, sandy or rocky dolomitic outcrops; Dms, dwarf mountain scrub dominated by cushion plants; For-Cl, forest clearings and edges; For-In, forest interior; Gra, grasslands and meadows; Scle, Tall Mediterranean sclerophyllous forest and scrub; Str, Banks of permanent streams or flooded areas surrounding springs.
