## Appendix S2 for "Unclusterable, underdispersed arrangement of insect-pollinated plants in pollinator niche space"

**Supporting Information.** Carlos M. Herrera. Unclusterable, underdispersed arrangement of insect-pollinated plants in pollinator niche space. *Ecology*.

**Appendix S2 Number of taxonomic categories of insect pollinators recorded per plant species.**

TABLE S1. Number of different orders, families, genera and species of insect pollinators recorded for each of the plant species studied ($N$ = 221). Figures refer to pollinators that were reliably identified up to the corresponding level of taxonomic resolution, which explains why sometimes Species < Genera.

| Species | Orders | Families | Genera | Species |
|---|---|---|---|---|
| *Achillea odorata* | 4 | 26 | 32 | 26 |
| *Acinos alpinus* | 3 | 12 | 16 | 20 |
| *Allium roseum* | 5 | 15 | 26 | 25 |
| *Allium scorodoprasum* | 4 | 24 | 61 | 66 |
| *Allium sphaerocephalon* | 4 | 15 | 29 | 34 |
| *Alyssum simplex* | 4 | 5 | 5 | 5 |
| *Amelanchier ovalis* | 3 | 11 | 16 | 19 |
| *Anagallis monelli* | 3 | 7 | 9 | 10 |
| *Anarrhinum laxiflorum* | 3 | 4 | 4 | 7 |
| *Andryala ragusina* | 4 | 18 | 27 | 25 |
| *Anthemis pedunculata* | 4 | 16 | 18 | 16 |
| *Anthericum liliago* | 4 | 7 | 8 | 8 |
| *Anthyllis vulneraria* | 3 | 5 | 7 | 9 |
| *Antirrhinum australe* | 1 | 1 | 2 | 2 |
| *Aphyllanthes monspeliensis* | 5 | 17 | 42 | 38 |
| *Aquilegia cazorlensis* | 4 | 6 | 7 | 12 |
| *Aquilegia vulgaris* | 2 | 5 | 9 | 11 |
| *Arbutus unedo* | 2 | 3 | 4 | 5 |
| *Arenaria armerina* | 3 | 14 | 19 | 16 |

| | | | | |
|---|---|---|---|---|
| *Arenaria modesta* | 2 | 4 | 5 | 6 |
| *Arenaria obtusiflora* | 3 | 19 | 21 | 14 |
| *Armeria filicaulis* | 6 | 22 | 22 | 21 |
| *Armeria villosa* | 4 | 20 | 20 | 21 |
| *Asphodelus cerasiferus* | 4 | 12 | 15 | 22 |
| *Astragalus bourgaeanus* | 1 | 3 | 3 | 5 |
| *Astragalus incanus* | 2 | 2 | 3 | 6 |
| *Atropa baetica* | 2 | 4 | 5 | 6 |
| *Bellis perennis* | 5 | 16 | 15 | 12 |
| *Bellis sylvestris* | 4 | 13 | 12 | 11 |
| *Berberis hispanica* | 3 | 17 | 23 | 26 |
| *Biscutella laxa* | 4 | 11 | 12 | 7 |
| *Calamintha nepeta* | 3 | 8 | 11 | 8 |
| *Campanula dieckii* | 4 | 14 | 17 | 16 |
| *Campanula mollis* | 3 | 5 | 5 | 4 |
| *Carduncellus monspelliensium* | 4 | 17 | 27 | 21 |
| *Carduus platypus* | 3 | 10 | 17 | 17 |
| *Carduus tenuiflorus* | 4 | 14 | 20 | 18 |
| *Carlina hispanica* | 4 | 13 | 37 | 34 |
| *Carlina racemosa* | 4 | 7 | 21 | 17 |
| *Catananche caerulea* | 4 | 17 | 32 | 30 |
| *Centaurea calcitrapa* | 4 | 10 | 30 | 35 |
| *Centaurea graminifolia* | 4 | 13 | 21 | 19 |
| *Cerastium gibraltaricum* | 4 | 11 | 12 | 11 |
| *Chaenorhinum macropodum* | 3 | 4 | 4 | 6 |
| *Chiliadenus glutinosus* | 4 | 12 | 23 | 21 |
| *Chondrilla juncea* | 4 | 13 | 27 | 22 |
| *Cirsium acaule* | 4 | 13 | 27 | 26 |
| *Cirsium monspessulanum* | 4 | 14 | 30 | 27 |

| | | | | |
|---|---|---|---|---|
| *Cirsium pyrenaicum* | 3 | 14 | 38 | 42 |
| *Cirsium vulgare* | 3 | 9 | 17 | 18 |
| *Cistus albidus* | 3 | 15 | 22 | 19 |
| *Cistus monspeliensis* | 4 | 26 | 44 | 39 |
| *Cistus salviifolius* | 3 | 14 | 21 | 21 |
| *Cleonia lusitanica* | 3 | 9 | 16 | 12 |
| *Clinopodium vulgare* | 3 | 6 | 10 | 9 |
| *Conopodium arvense* | 3 | 12 | 12 | 10 |
| *Convolvulus arvensis* | 4 | 14 | 20 | 18 |
| *Coris monspeliensis* | 5 | 9 | 14 | 13 |
| *Crataegus monogyna* | 3 | 14 | 20 | 23 |
| *Crepis albida* | 4 | 7 | 7 | 6 |
| *Crocus nevadensis* | 2 | 3 | 3 | 4 |
| *Crocus nudiflorus* | 3 | 5 | 8 | 8 |
| *Cuscuta triumvirati* | 4 | 17 | 26 | 19 |
| *Daphne gnidium* | 3 | 9 | 15 | 14 |
| *Daphne laureola* | 4 | 6 | 7 | 10 |
| *Digitalis obscura* | 1 | 5 | 8 | 9 |
| *Draba hispanica* | 4 | 11 | 11 | 9 |
| *Echinospartum boissieri* | 2 | 5 | 8 | 11 |
| *Echium flavum* | 3 | 7 | 13 | 18 |
| *Erinacea anthyllis* | 3 | 7 | 11 | 17 |
| *Erodium cazorlanum* | 4 | 9 | 7 | 7 |
| *Erodium cheilanthifolium* | 4 | 11 | 12 | 12 |
| *Erodium cicutarium* | 3 | 11 | 11 | 7 |
| *Erophila verna* | 2 | 4 | 3 | 6 |
| *Eryngium campestre* | 4 | 32 | 70 | 69 |
| *Eryngium dilatatum* | 4 | 22 | 49 | 50 |
| *Erysimum cazorlense* | 5 | 9 | 9 | 9 |
| *Erysimum medio-hispanicum* | 4 | 14 | 15 | 13 |

| | | | | |
|---|---|---|---|---|
| *Euphorbia nicaeensis* | 4 | 32 | 52 | 45 |
| *Filipendula vulgaris* | 5 | 14 | 17 | 14 |
| *Fumana baetica* | 4 | 13 | 23 | 22 |
| *Fumana paradoxa* | 4 | 11 | 16 | 17 |
| *Fumana procumbens* | 3 | 8 | 9 | 6 |
| *Gagea soleirolii* | 4 | 13 | 15 | 13 |
| *Galatella linosyris* | 4 | 18 | 34 | 29 |
| *Galium verum* | 5 | 20 | 30 | 29 |
| *Genista longipes* | 2 | 5 | 5 | 6 |
| *Genista pseudopilosa* | 1 | 4 | 4 | 6 |
| *Genista scorpius* | 2 | 4 | 6 | 8 |
| *Geranium molle* | 4 | 10 | 11 | 11 |
| *Geum sylvaticum* | 4 | 12 | 17 | 16 |
| *Gladiolus illyricus* | 3 | 13 | 24 | 25 |
| *Globularia spinosa* | 6 | 13 | 12 | 8 |
| *Gymnadenia conopsea* | 1 | 3 | 3 | 5 |
| *Hedera helix* | 3 | 15 | 24 | 24 |
| *Helianthemum apenninum* | 5 | 21 | 30 | 25 |
| *Helianthemum cinereum* | 3 | 7 | 8 | 3 |
| *Helianthemum oelandicum* | 4 | 12 | 17 | 15 |
| *Helleborus foetidus* | 1 | 4 | 6 | 5 |
| *Hepatica nobilis* | 2 | 3 | 2 | 4 |
| *Himantoglossum hircinum* | 2 | 5 | 7 | 6 |
| *Hormathophylla baetica* | 3 | 5 | 5 | 5 |
| *Hormathophylla spinosa* | 3 | 8 | 9 | 8 |
| *Hyacinthoides reverchonii* | 2 | 7 | 8 | 8 |
| *Hypericum caprifolium* | 3 | 8 | 11 | 11 |
| *Hypericum ericoides* | 3 | 4 | 9 | 7 |
| *Hypericum perforatum* | 5 | 20 | 33 | 42 |
| *Iberis carnosa* | 2 | 3 | 4 | 4 |

| | | | | |
|---|---|---|---|---|
| *Inula montana* | 4 | 21 | 48 | 44 |
| *Iris foetidissima* | 1 | 1 | 2 | 4 |
| *Iris planifolia* | 1 | 3 | 3 | 4 |
| *Jasonia tuberosa* | 4 | 11 | 20 | 19 |
| *Jonopsidium prolongoi* | 5 | 11 | 9 | 7 |
| *Jurinea humilis* | 5 | 13 | 16 | 14 |
| *Klasea nudicaulis* | 5 | 20 | 24 | 25 |
| *Klasea pinnatifida* | 4 | 20 | 44 | 46 |
| *Knautia subscaposa* | 4 | 22 | 50 | 52 |
| *Lamium amplexicaule* | 1 | 3 | 4 | 6 |
| *Lavandula latifolia* | 3 | 19 | 52 | 63 |
| *Leontodon longirrostris* | 3 | 11 | 14 | 10 |
| *Leucanthemopsis pallida* | 4 | 14 | 14 | 12 |
| *Linaria aeruginea* | 2 | 3 | 3 | 3 |
| *Linaria verticillata* | 1 | 2 | 3 | 4 |
| *Linaria viscosa* | 2 | 3 | 6 | 8 |
| *Linum bienne* | 3 | 4 | 4 | 6 |
| *Linum narbonense* | 4 | 8 | 9 | 6 |
| *Linum tenue* | 4 | 16 | 28 | 29 |
| *Lithodora fruticosa* | 3 | 7 | 11 | 16 |
| *Lonicera arborea* | 2 | 4 | 6 | 9 |
| *Lysimachia ephemerum* | 4 | 20 | 34 | 37 |
| *Lythrum junceum* | 4 | 8 | 9 | 7 |
| *Lythrum salicaria* | 3 | 6 | 13 | 15 |
| *Mantisalca salmantica* | 4 | 25 | 48 | 59 |
| *Marrubium supinum* | 4 | 16 | 27 | 27 |
| *Mentha aquatica* | 4 | 13 | 18 | 19 |
| *Merendera montana* | 3 | 10 | 16 | 15 |
| *Narcissus bujei* | 3 | 6 | 7 | 12 |
| *Narcissus cuatrecasasii* | 3 | 6 | 7 | 10 |

| | | | | |
|---|---|---|---|---|
| *Narcissus hedraeanthus* | 3 | 5 | 5 | 5 |
| *Narcissus longispathus* | 4 | 12 | 18 | 26 |
| *Narcissus triandrus* | 1 | 2 | 2 | 4 |
| *Nepeta tuberosa* | 3 | 6 | 8 | 8 |
| *Ononis pusilla* | 1 | 3 | 4 | 4 |
| *Ononis spinosa* | 1 | 2 | 5 | 8 |
| *Orchis coriophora* | 3 | 14 | 19 | 18 |
| *Origanum virens* | 3 | 11 | 13 | 13 |
| *Ornithogalum umbellatum* | 5 | 14 | 14 | 11 |
| *Orobanche haenseleri* | 1 | 2 | 3 | 3 |
| *Papaver dubium* | 4 | 7 | 12 | 13 |
| *Parnassia palustris* | 2 | 3 | 3 | 3 |
| *Petrorhagia nanteuilii* | 4 | 11 | 18 | 13 |
| *Phlomis herba-venti* | 3 | 9 | 18 | 15 |
| *Phlomis lychnitis* | 2 | 4 | 7 | 6 |
| *Picnomon acarna* | 3 | 8 | 18 | 23 |
| *Pilosella pseudopilosella* | 4 | 19 | 24 | 18 |
| *Pistorinia hispanica* | 3 | 6 | 10 | 9 |
| *Plumbago europaea* | 3 | 12 | 32 | 34 |
| *Polygala boissieri* | 3 | 5 | 6 | 5 |
| *Polygonatum odoratum* | 1 | 1 | 2 | 3 |
| *Potentilla caulescens* | 4 | 10 | 17 | 18 |
| *Potentilla reptans* | 3 | 12 | 12 | 10 |
| *Primula acaulis* | 3 | 4 | 4 | 7 |
| *Prolongoa hispanica* | 5 | 12 | 16 | 10 |
| *Prunus mahaleb* | 3 | 10 | 14 | 17 |
| *Pterocephalus spathulatus* | 3 | 10 | 21 | 21 |
| *Ptilostemon hispanicus* | 3 | 11 | 26 | 27 |
| *Ranunculus bulbosus* | 5 | 19 | 23 | 17 |
| *Ranunculus ficaria* | 3 | 8 | 8 | 8 |

| | | | | |
|---|---|---|---|---|
| *Ranunculus malessanus* | 4 | 16 | 16 | 14 |
| *Ranunculus paludosus* | 4 | 17 | 22 | 21 |
| *Ranunculus repens* | 4 | 18 | 28 | 18 |
| *Rhagadiolus edulis* | 3 | 10 | 10 | 7 |
| *Roemeria argemone* | 3 | 5 | 4 | 4 |
| *Rosa canina* | 3 | 12 | 18 | 19 |
| *Rosa micrantha* | 4 | 20 | 41 | 43 |
| *Rosa sicula* | 3 | 8 | 12 | 13 |
| *Rosmarinus officinalis* | 3 | 13 | 22 | 28 |
| *Rubus ulmifolius* | 4 | 18 | 44 | 46 |
| *Salvia lavandulifolia* | 4 | 8 | 10 | 9 |
| *Salvia verbenaca* | 2 | 5 | 9 | 6 |
| *Santolina rosmarinifolia* | 5 | 28 | 42 | 38 |
| *Saponaria ocymoides* | 2 | 3 | 4 | 7 |
| *Satureja intricata* | 4 | 23 | 43 | 45 |
| *Saxifraga haenseleri* | 3 | 9 | 9 | 5 |
| *Saxifraga carpetana* | 3 | 8 | 6 | 5 |
| *Scabiosa andryaefolia* | 3 | 11 | 26 | 27 |
| *Scilla paui* | 2 | 6 | 7 | 8 |
| *Scorzonera albicans* | 5 | 8 | 8 | 6 |
| *Sedum album* | 5 | 30 | 57 | 52 |
| *Sedum dasyphyllum* | 3 | 6 | 10 | 7 |
| *Sedum mucizonia* | 4 | 8 | 11 | 11 |
| *Sedum sediforme* | 4 | 14 | 21 | 24 |
| *Senecio doria* | 4 | 14 | 20 | 20 |
| *Senecio malacitanus* | 3 | 13 | 22 | 22 |
| *Seseli montanum* | 3 | 11 | 19 | 12 |
| *Sideritis incana* | 3 | 11 | 18 | 14 |
| *Silene colorata* | 3 | 6 | 6 | 8 |
| *Silene psammitis* | 3 | 7 | 10 | 7 |

| | | | | |
|---|---|---|---|---|
| *Sisymbrella aspera* | 4 | 18 | 21 | 20 |
| *Sisymbrium crassifolium* | 3 | 8 | 8 | 7 |
| *Solidago virgaurea* | 2 | 5 | 8 | 7 |
| *Sonchus aquatilis* | 4 | 11 | 22 | 20 |
| *Stachys officinalis* | 3 | 15 | 27 | 27 |
| *Taraxacum laevigatum* | 5 | 14 | 20 | 18 |
| *Teucrium aureum* | 3 | 18 | 32 | 36 |
| *Teucrium rotundifolium* | 3 | 6 | 7 | 10 |
| *Teucrium webbianum* | 2 | 4 | 7 | 7 |
| *Thymus mastichina* | 4 | 17 | 32 | 32 |
| *Thymus orospedanus* | 3 | 14 | 21 | 17 |
| *Thymus serpylloides* | 3 | 8 | 15 | 14 |
| *Trifolium campestre* | 3 | 8 | 11 | 7 |
| *Ulex parviflorus* | 2 | 5 | 6 | 7 |
| *Urginea maritima* | 3 | 10 | 21 | 20 |
| *Valeriana tuberosa* | 4 | 14 | 22 | 20 |
| *Verbascum giganteum* | 3 | 9 | 18 | 17 |
| *Verbascum sinuatum* | 3 | 8 | 20 | 18 |
| *Viburnum tinus* | 3 | 14 | 17 | 13 |
| *Vicia onobrychioides* | 3 | 4 | 9 | 8 |
| *Vicia pseudocracca* | 3 | 4 | 8 | 11 |
| *Viola cazorlensis* | 2 | 2 | 2 | 3 |
| *Viola odorata* | 1 | 2 | 3 | 6 |
